# Cajal bodies shape the nuclear transcriptome, intron retention and transcripts redistribution in *Arabidopsis* roots under hypoxia

**DOI:** 10.64898/2026.10.02.756184

**Authors:** S Górka, D Kubiak, M Szcześniak, A Kołowerzo-Lubnau, A Zienkiewicz, D Dombek, Sz Kowalczyk, K Niedojadło, J Niedojadło

## Abstract

Cajal bodies (CBs) are nuclear condensates known for their role in sn- and snoRNA maturation in animal, but their function in plant stress responses remains poorly understood. We compared wild-type (WT) *Arabidopsis thaliana* with the CB-lacking *ncb-1* mutant during hypoxia and reoxygenation, focusing on nuclear retention of transcripts. In both genotypes, hypoxia caused nuclear accumulation of poly(A^+^) RNA and extensive reprogramming of gene expression. However, the transcriptional response, including induction of hypoxia-responsive genes, was considerably weaker in *ncb-1*, both during stress and recovery. In nuclear RNA, introns with low (intron retention index, IRX < 10%) and high (IRX > 50%) retention showed opposite, genotype-dependent dynamics. During hypoxia, efficiently spliced introns (IRX < 10%) accounted for over 70% of introns detected in nuclear RNA. Relative to normoxia, this proportion increased two- to threefold in WT but only moderately in the mutant. Together with poly(A^+^) RNA accumulation, this suggests that largely spliced transcripts, including those of genes involved in protein transport and ribosome biogenesis, are held in the nucleus during stress. Conversely, strongly retained introns, often terminal and located in genes related to hypoxia and other abiotic stresses, were most abundant in WT nuclear RNA under normoxia. Their proportion dropped sharply during hypoxia, and much less so in *ncb-1*. This indicate that WT plants maintain a nuclear reserve of incompletely spliced stress-related transcripts that are processed upon stress. The mutant also showed reduced interaction between U2 snRNA and U2B″, which leads to splicing defects. Our results indicate that CBs regulate which transcripts are held in the nucleus and how they are processed, contributing to plant tolerance of oxygen deprivation and recovery.

## Introduction

Cajal bodies (CBs) are membraneless nuclear bio-condensates whose assembly is thought to involve liquid-liquid phase separation (LLPS) and multivalent protein-protein and protein-RNA interactions in the cell nucleus (Riback et al., 2020; Hedtfeld et al., 2024). In eukaryotic cells, CBs are structurally and functionally defined by the presence of several major components, including coilin, a CB marker protein; fibrillarin, a methyltransferase associated with C/D box small nucleolar ribonucleoproteins (snoRNPs); and SmB/B′, core components of spliceosomal snRNPs involved in pre-mRNA splicing (Taliansky et al., 2023). In addition, WRAP53β, also known as WD40-encoding RNA antisense to p53, is essential for proper CB formation in animal models. WRAP53β stabilizes the interaction between coilin and survival motor neuron protein (SMN), which is involved in RNA splicing and the repair of DNA double-strand breaks (Mahmoudi et al., 2010; Bergstrand et al., 2019). In addition to the functions carried out via proteins, CBs are an important hub for the metabolism of different RNAs (Taliansky et al., 2023). Their canonical role includes the biogenesis of snRNA and snoRNA and the assembly of snRNP complexes, which is crucial for pre-mRNA splicing and rRNA maturation (Ohtani 2018; Basello et al., 2025). A unique feature of CBs in plants is their involvement in mRNA and siRNA metabolism (Li et al. 2006; Niedojadło et al., 2014; Górka et al., 2022). Recent studies have further refined the CBs as dynamic condensates rather than static structures defined solely by the presence of coilin, with RNA considered a key structural component of such nuclear membraneless bodies (André and Spruijt, 2020). In particular, the N-terminal domain of coilin has been shown to mediate multivalent interactions with Nopp140 in human cells, supporting bio-condensation and CB maintenance. Moreover, recent interactome-based approaches have identified additional coilin-associated factors and potential CB components, indicating that the molecular composition of CBs is broader and more dynamic than previously assumed. These findings support the that CBs are not merely passive sites of snRNP maturation but highly regulated nuclear hubs that integrate RNA metabolism, ribonucleoprotein assembly, and stress-responsive nuclear organization (Courchaine et al., 2022; Arias Escayola et al., 2025).

Recent studies have shown that CBs are involved in plant responses to both abiotic and biotic stresses (Shaw et al., 2014; Niedojadło et al., 2014, 2016; Love et al., 2017). During viral infection, coilin and fibrillarin can participate in the formation and trafficking of cytoplasmic viral ribonucleoprotein complexes that move systemically through the phloem (Kim et al., 2007; Canetta et al., 2008). It has been shown that the interaction of the NIa (nuclear inclusion protein) protein encoded by Potato virus A (PVA)with fibrillarin can target components of the RNA-silencing machinery, thereby suppressing RNA silencing and facilitating viral infection (Rajamäki and Valkonen 2009). However, there are also examples in which coilin restricts viral infection. Viral proteins can interact with coilin and relocate it from CBs to the nucleolus, enhancing the accumulation of salicylic acid (SA), which stimulates downstream components of the SA-mediated defense pathway (Shaw et al., 2019). Our studies have shown that CBs also play an important role in plant responses to abiotic stress. The *ncb-1* mutant, which lacks functional CBs, displays reduced tolerance to hypoxic stress and impaired recovery following reoxygenation (Niedojadło et al., 2016; Górka et al., 2022). This may be associated with the involvement of CBs in nuclear RNA retention (Niedojadło et al., 2014). Nuclear accumulation of poly(A)+ RNA during hypoxia is reduced in plants lacking Cajal bodies, both in leaves and roots (Niedojadło et al., 2016; Górka et al., 2022). However, the role of CBs in this process is not clear. In our other studies, we demonstrated that in *ncb-1* mutants, intron retention was observed in the mRNA encoding RPB1, the largest subunit of RNA polymerase II (Górka et al., 2022). The intron-retaining RPB1 isoform accumulates at high levels in the nucleus. These observations suggest that intron retention itself may contribute to nuclear RNA retention, independently of or in addition to CB-mediated sequestration. Nuclear accumulation of intron-containing mRNAs may allow these transcripts to evade degradation by nonsense-mediated mRNA decay (NMD) (Kalyna et al., 2012; Göhring et al., 2014). During reoxygenation, retained introns may be removed posttranscriptionally from incompletely spliced transcripts that accumulated in the nuclei of *ncb-1* plants during hypoxia. Nevertheless, the role of CBs in plant stress responses is not fully understood and requires further research (Love et al., 2017; Makhotenko et al., 2019).

Oxygen deprivation (hypoxia) occurs in plant roots during flooding and waterlogging and forces a rapid shift from mitochondrial respiration to anaerobic metabolism. In *Arabidopsis*, a set of approximately 50 core hypoxia-responsive genes, including those encoding alcohol dehydrogenase (*ADH1*), pyruvate decarboxylase (*PDC1*), sucrose synthases and hypoxia-responsive universal stress proteins, is induced in virtually all cell types (Mustroph et al., 2009). Their expression is controlled largely by group VII ethylene response factors, which are stabilized under low oxygen through the oxygen-dependent N-end rule pathway (Gibbs et al., 2011; Licausi et al., 2011). At the same time, hypoxia strongly restricts global translation: only a subset of mRNAs, mostly those encoding stress-related proteins, remains associated with polysomes, whereas many other transcripts are stored and rapidly re-engaged in translation after reoxygenation (Branco-Price et al., 2008). Integrative analyses have further shown that the response to transient hypoxia in *Arabidopsis* is regulated predominantly at the nuclear level, from chromatin accessibility to nuclear transcript accumulation (Lee and Bailey-Serres 2019). Post-transcriptional control of mRNA processing and localization is therefore an integral part of the hypoxia response.

Introns are important regulatory elements that can influence gene expression at multiple levels, and their effects depend on their sequence, position within the gene, and efficiency of recognition by the splicing machinery. The presence of introns can enhance gene expression through intron-mediated enhancement (IME), which has been associated with increased transcriptional activity and mRNA accumulation (Parra et al., 2011; Dwyer et al., 2021; Kowal et al., 2025). In addition, intronic sequences contain cis-regulatory elements that influence splice-site recognition and spliceosome assembly and can therefore promote or repress the efficiency of pre-mRNA splicing (Reddy et al., 2012). Retention of an intron further expands the regulatory potential of a transcript by altering its sequence and coding capacity. Retained introns may introduce additional or alternative open reading frames, upstream open reading frames (uORFs), premature termination codons (PTCs), or other regulatory elements, thereby affecting transcript stability, localization, and translational efficiency (Ner-Gaon et al., 2004; Jacob and Smith 2017; Martín et al., 2021). One important consequence of intron retention is its connection with nonsense-mediated mRNA decay (NMD). When intron retention introduces a PTC located more than approximately 50–55 nucleotides upstream of a downstream exon–exon junction, the transcript can be recognized by the EJC-dependent NMD pathway (Singh et al., 2008; Bicknell et al., 2012; Karousis and Mühlemann 2022; Benslimane et al., 2024). NMD acts not only as an mRNA quality-control mechanism that prevents the accumulation and translation of aberrant PTC-containing transcripts but also as an important component of post-transcriptional gene regulation. In plants, NMD modulates the abundance of physiological transcripts involved in development, hormonal signalling, circadian rhythms, reproduction, pathogen defence, and responses to environmental stress (Raxwal and Riha 2023; Luha et al., 2024). However, intron-retaining transcripts are not invariably targeted by NMD. Some remain temporarily retained in the nucleus and may subsequently undergo post-transcriptional splicing, providing a reservoir of transcripts that can be rapidly mobilized in response to developmental or environmental signals (Boothby et al., 2013; Göhring et al., 2014; Niedojadło et al., 2016; Jia et al., 2020; Rudzka et al., 2022).

Despite this growing body of evidence, how Cajal bodies shape the hypoxic transcriptome and splicing landscape at a genome-wide scale remains largely unexplored. Here, we combine microscopic, transcriptomic, and splicing analyses of wild-type and *ncb-1 Arabidopsis* plants to dissect how these nuclear condensates coordinate RNA retention in nucleus, gene expression reprogramming, and intron processing during hypoxia and subsequent recovery. Our findings extend the previously proposed role of Cajal bodies in the hypoxia response, providing a more detailed molecular picture of how these structures contribute to plant resilience under oxygen deprivation.

## Materials and methods

### Hydroponic cultivation

Arabidopsis thaliana (Col-0) wild-type plants expressing U2B″–GFP (WT) and the ncb-1 mutant, which was isolated in the U2B″–GFP background (Collier et al. 2006), were grown hydroponically in an Araponics system using FloraSeries/TriPart nutrient solutions prepared at a concentration of 0.7 mL/L of each component, as described by Tocquin et al. (2003). After brief soaking in distilled water, seeds were sown at a density of three per well in 0.65% agar (Sigma-Aldrich). Seedlings were cultivated for 2 weeks at 22°C under a 16 h light/8 h dark photoperiod and a light intensity of 120–150 µmol m⁻² s⁻¹. Aeration of the nutrient solutions was initiated after 1 week. Two-week-old seedlings were subjected to 5 days of hypoxic stressby submerge in water under limited light conditions. After the stress treatment, roots were collected and frozen, while the remaining plants were subjected to 1 day of reoxygenation, after which the material was collected again. All root samples were stored at −80°C.

### Isolation of nuclei (INTACT) and nuclear RNA

To enable affinity purification of nuclei from the wt and *ncb-1* background, *ncb-1* plants were crossed with the INTACT line *pUBQ10:NTF/pACT2:BirA* (Col-0 background) Lee and Bailey-Serres, 2019). Roots (0.8 g) were ground in liquid nitrogen and suspended in 6 mL of NPB buffer (0× Nuclei Purification Buffer (10× NPB): 200 mM MOPS (pH 7.0), 400 mM NaCl, 900 mM KCl, 20 mM EDTA, 5 mM EGTA). The resulting suspension was filtered through a nylon sieve with a mesh size of 30 µm. The filtrate was centrifuged for 7 min at 1,200 × g and 4°C, after which the nuclei-containing pellet was resuspended in 1 mL of NPB. The nuclear suspension was incubated with 25 μL of pre-washed Dynabeads M-280 Streptavidin for 40 min on a rotator at 4°C. The samples were then washed twice with 13 mL of NPBt, with each wash followed by magnetic separation for 5 min. Finally, the pellet was resuspended in 20 μL of NPB (Lee and Bailey-Serres 2019).

RNA was isolated using the RNeasy Micro Kit (Qiagen). The nuclear suspension was lysed in 350 μL of RLT buffer containing 3.5 μL of 2-mercaptoethanol, vortexed for 1.5 min, and incubated for 5 min at room temperature. Following centrifugation, 340 μL of 70% ethanol was added to the collected supernatant, and the resulting mixture was loaded onto an RNeasy MinElute column. The column was sequentially washed with 700 μL of RW1 buffer, 500 μL of RPE buffer, and 500 μL of 80% ethanol. After the addition of ethanol, the column was centrifuged for 2 min at 10,000 rpm and subsequently dried by centrifugation for 5 min at maximum speed. RNA was eluted in 14 μL of RNase-free water.

### Total RNA isolation

Total RNA was isolated from the roots of Arabidopsis thaliana using the Direct-zol™ RNA MiniPrep Plus kit (Zymo Research, USA), according to the manufacturer’s protocol. Briefly, 1000 µL of TRI Reagent (Sigma-Aldrich) was added to 100 mg of ground tissue. During the isolation procedure, on-column DNA digestion was performed using DNase I (1500 U). RNA was washed according to the kit protocol and eluted in DNase/RNase-free water. After isolation, the samples were stored at −80°C until further analysis. The concentration and purity of the isolated RNA were initially assessed spectrophotometrically using a NanoDrop One spectrophotometer (Thermo Scientific) based on the A260/A280 and A260/A230 absorbance ratios. Then RNA integrity was evaluated by capillary electrophoresis using the RNA 6000 Nano Assay Kit and the Total RNA Eukaryotic Pico Kit with the 2100 Bioanalyzer system (Agilent Technologies), according to the manufacturer’s instructions. RNA integrity was expressed as the RNA integrity number (RIN), and only samples with RIN values above 6 were selected for further analyses (Schroeder A. et al. 2006).

### Library preparation and sequence analysis

cDNA libraries were prepared from nuclear and total RNA isolated from root cells of Arabidopsis thaliana wild-type plants and the *ncb-1* mutant, derived from plants grown under control conditions, subjected to 5 days of hypoxia, and 1 day of reoxygenation following hypoxia. After RNA quality and quantity assessment, rRNA depletion was performed using the RiboMinus™ Plant Kit (Invitrogen), and the RNA was subsequently concentrated according to the manufacturer’s protocol. After RQ1 Rnase-Free DNase (Promega) treatment cDNA libraries were prepared using the NEBNext® Ultra™ II for Illumina kit (New England BioLabs). Three biological replicates were analysed for each genotype, condition and RNA fraction (36 libraries in total). Following library preparation, paired-end sequencing (2 × 100 nt) was performed by Fasteris SA (Plan-les-Ouates, Switzerland) on the Illumina NovaSeq SP platform using the NVS-SP-2× sequencing method.

### Fluorescence in situ hybridization (FISH) and confocal microscopy

For fluorescence and confocal microscopy, roots were cut into small fragments and fixed in 4% (w/v) paraformaldehyde (PFA) in 0.1 M PBS, pH 7.2, for 12 h. Before embedding in BMM resin, samples were dehydrated through a graded ethanol series (10–96%, 10 min each). Embedded material was sectioned on a Leica UCT ultramicrotome to obtain sections (1.5 µm), which were mounted on BioBond-coated slides (British Biocell International). Before hybridization, the BMM resin was removed by washing the sections in pure acetone twice for 15 min and then for 5 min in 4× SSC (pH 7.0). The FISH reaction was preceded by 1 h pre-hybridization in hybridization buffer containing 50% (v/v) hybridization buffer (Merck), 30% (v/v) formamide and 20% (v/v) water. This was followed by overnight (≥12 h) in situ hybridization with specific probes 5′-end–labeled with Cy3 (dilution 1:250). After hybridization, sections were washed in 4×, 2× and 1× SSC. Nuclei were counterstained with Dapi (Merck) at 1:1000 in 2× SSC for 10 min. Negative controls were processed identically, except that the probes were omitted.

### RNA immunoprecipitation (RIP) and RT-qPCR

RNA immunoprecipitation (RIP) was performed using roots of *Arabidopsis thalian*a from the WT and *ncb-1* lines expressing U2B″–GFP. Approximately 0.8 g of frozen root tissue was ground in liquid nitrogen and resuspended in 1.2 mL of lysis buffer (100 mM Tris–HCl, pH 7.5; 10% sucrose; 5 mM EDTA; 300 mM NaCl; 0.15% SDS; 1% Triton X-100; protease inhibitor, 1:100; RNaseOUT Recombinant Ribonuclease Inhibitor, 1 U µL⁻¹) prepared in DEPC-treated water.

The lysate was vortexed, incubated on ice, and centrifuged at 14,000 rpm for 5 min at 4 °C. The clarified supernatant was transferred to a chilled tube and diluted with dilution buffer (10 mM Tris–HCl, pH 7.5; 150 mM NaCl; 0.5 mM EDTA) at a ratio of 4:6 (lysate:buffer) supplemented with inhibitors. A 25 µL aliquot was collected as the input control (Input I).

GFP-tagged protein complexes were immunoprecipitated using GFP-Trap magnetic particles (Chromotek), which were incubated with diluted lysates on a rotator (3–4 rpm) for 24 h at 4 °C. After incubation, the beads were magnetically separated and 50 µL of the supernatant was collected as a second input control (Input II). The beads were then washed sequentially with low-salt buffer (20 mM Tris–HCl, pH 8.0; 2 mM EDTA; 1% Triton X-100; 150 mM NaCl; 0.1% SDS), high-salt buffer (20 mM Tris–HCl, pH 8.0; 2 mM EDTA; 1% Triton X-100; 500 mM NaCl; 0.1% SDS), lithium chloride buffer (10 mM Tris–HCl, pH 8.0; 20 mM EDTA; 250 mM LiCl; 1% NP-40; 1% sodium deoxycholate), and TE buffer (10 mM Tris–HCl, pH 8.0; 10 mM EDTA) to remove nonspecifically bound molecules.

For RNA isolation, the beads were resuspended in RIP elution buffer (50 mM Tris–HCl, pH 6.8; 1 mM EDTA; 1% SDS) and treated with proteinase K (1 U per 40 µL), followed by incubation at 55 °C for 30 min and 60 °C for 30 min to release RNA from protein complexes. RNA was extracted using TRI Reagent, followed by chloroform phase separation and precipitation with isopropanol in the presence of sodium acetate and GlycoBlue coprecipitant. The RNA pellet was washed with 75% ethanol, air-dried, and resuspended in RNase-free water.

Residual DNA was removed using RQ1 RNase-free DNase (Promega). First-strand cDNA synthesis was performed using random hexamer primers (Thermo Fisher Scientific). Quantitative PCR (qPCR) was carried out to assess the enrichment of selected transcripts in RIP samples relative to the input controls.

The following primer pairs were used for qPCR: AT-U2-2RF consisting of AT-U2-2F (5′-ACC CGC CAA ACT AGA AGT GA-3′; Tm = 51.8 °C) and AT-U2-2R (5′-AGA CAC ACA TGA CGC AGA AGA-3′; Tm = 51.8 °C), and AT-U2-3RF consisting of AT-U2-3F (5′-AGA CAA CAC CCC AAG AGC TT-3′; Tm = 51.8 °C) and AT-U2-3R (5′-CGA CTG GAG CTA TGC AAT CG-3′; Tm = 53.8 °C). Real-time PCR was performed using a LightCycler 480 SYBR Green Master Kit (Roche). The sequences of the primers used are shown in Supplementary Table S1. The thermal profile was carried out according to default parameters of a pre-incubation step at 95 °C for 10 min (1 cycle) and an amplification step at 95 °C for 10 s, 52 °C for 10 s, and 72 °C for 20 s (45 cycles). RIP-qPCR data were analysed using the percent input method. Cq values for the 1% input samples were corrected for the input dilution, and RNA enrichment in the immunoprecipitated fractions was expressed as the percentage of input RNA (Fuentes-Iglesias et al. 2020).

### Western blot analysis

For Western blot analysis, beads obtained after immunoprecipitation (see above) and suspended in TE buffer were mixed with 2× SDS Laemmli sample buffer (150 mM Tris–HCl, pH 6.8; 10% β-mercaptoethanol; 1.2% SDS; 30% glycerol; 0.4% bromophenol blue) and incubated at 95 °C for 10 min to elute and denature proteins.

Proteins were separated on a 12% SDS–polyacrylamide gel prepared using the TGX FastCast Acrylamide Kit (Bio-Rad) and run in a Mini-PROTEAN 3 electrophoresis system (Bio-Rad, USA). Proteins were transferred onto PVDF membranes (Merck), which were blocked for 1 h in TBST [20 mM Tris-HCl, pH 7.5, and 150 mM NaCl and 0.05% (v/v) Tween 20] containing 5% (w/v) nonfat dry milk. Immunobloting was performed using anti-GFP:HRP antibody (Miltenyi Biotec) diluted 1:1000 in TBST for 12 h at 4 °C. Signals were detected using ECL Prime Western Blotting Detection Reagent (Cytiva) and visualized with a ChemiDoc MP imaging system (Bio-Rad).

### RNA-seq data processing

Raw paired-end reads were quality- and adapter-trimmed with BBDuk (BBTools), and reads derived from ribosomal RNA were removed by alignment with Bowtie2. For transcriptome assembly and intron-retention analysis, the clean reads were aligned to the *Arabidopsis thaliana* genome (TAIR10) with STAR (Dobin et al. 2013) in two-pass mode (parameters in “Read alignment for intron retention analysis”). A reference transcriptome was assembled with StringTie (Pertea et al. 2015) and used to quantify transcript- and gene-level expression with RSEM (Li and Dewey 2011) and Bowtie2 (Langmead and Salzberg 2012).

### Differential gene expression analysis

Differential expression was tested with DESeq2 (Love et al. 2014) using RSEM expected counts as input, separately for the nuclear and total RNA fractions. Genes with a Benjamini–Hochberg-adjusted p-value < 0.05 were considered differentially expressed. When the responses of the two genotypes were compared, genes were classified as differentially expressed in both genotypes (adjusted p < 0.05 in WT and in *ncb-1*) or exclusively in one genotype (adjusted p < 0.05 in that genotype only; Supplementary Table S1). Three within-genotype comparisons were performed for each genotype (hypoxia versus control, reoxygenation versus control, and reoxygenation versus hypoxia), together with three cross-genotype comparisons (*ncb-1* versus wild type under control, hypoxia and reoxygenation), giving nine comparisons per RNA fraction. Sample relationships were examined by principal component analysis (PCA) of regularized-log (rlog) expression of the 500 most variable genes; one nuclear RNA library (WT, hypoxia, replicate 2), which clustered with reoxygenation samples, was excluded from the differential expression analysis. Log₂ fold changes of core hypoxia-responsive genes (Mustroph et al. 2009) were compared between genotypes with the Wilcoxon signed-rank test. For the heatmap in Fig. 3E, genes differentially expressed in the same direction in both genotypes were ranked by the lower of the adjusted p-values from the hypoxia-versus-control and reoxygenation-versus-control comparisons, and row-wise z-scores of normalized nuclear RNA expression were displayed with pheatmap.

### Gene Ontology enrichment analysis

Gene Ontology (GO) over-representation analysis was performed with clusterProfiler (Wu et al. 2021) and the org.At.tair.db annotation database, separately for up-regulated (log₂FC > 0, adjusted p < 0.05) and down-regulated (log₂FC < 0, adjusted p < 0.05) gene sets, for the Biological Process (BP) and Cellular Component (CC) ontologies. P-values were adjusted by the Benjamini–Hochberg method, and terms with adjusted p < 0.05 were considered enriched. All annotated *A. thaliana* genes in org.At.tair.db were used as the background. The same procedure was applied to genes containing introns with IRX < 10% or > 50% (Figs. S5, S7).

### Cross-genotype comparison of fold changes

To compare the magnitude and direction of regulation between genotypes, log₂ fold changes from matched DESeq2 analyses (for example hypoxia versus control in wild type and in *ncb-1*) were plotted against one another. Genes were assigned to quadrants according to the sign of the fold change in each genotype and coloured by genotype-specific significance (adjusted p < 0.05 in one or both genotypes). GO enrichment was then performed separately for the genes in each quadrant and significance class, enabling identification of functional categories with genotype-dependent regulation.

### Comparison of total and nuclear RNA expression and identification of compartment switches

Compartment-specific expression was compared for each genotype and condition by plotting log₁₀(mean total RNA expression + 1) against log₁₀(mean nuclear RNA expression + 1) using FPKM values. A LOESS regression (span = 0.5) was fitted to model the expected nuclear–total relationship across all genes, and residuals were standardized by their standard deviation. Genes that reversed their compartment enrichment between conditions (“switchers”) were identified by comparing standardized LOESS residuals between two conditions (control versus hypoxia, or hypoxia versus reoxygenation). A gene was classified as a switcher when its standardized residual exceeded +1.5 in one condition and fell below −1.5 in the other (true sign reversal), and switchers were ranked by the magnitude of the change in standardized residual.

### snRNA and snoRNA expression analysis

Expression of small nuclear RNA (snRNA) and small nucleolar RNA (snoRNA) genes was quantified from the per-sample TPM matrix, with gene identity and RNA-type classification taken from the Ensembl annotation. For each condition, wild type and *ncb-1* were compared using two groups of three biological replicates. Genes were excluded if mean expression was zero in both genotypes or if fewer than three replicates were non-zero in both groups (at least one group was required to have three non-zero replicates). For each retained gene, the ratio of the higher to the lower group mean was computed, and genes with a ratio > 1.5 were considered differentially expressed between genotypes. Data were displayed as scatter plots of log₁₀-scaled expression (wild type versus *ncb-1*), coloured by RNA type.

### Read alignment for intron retention analysis

In parallel with RSEM quantification, reads were aligned to the *A. thaliana* genome (TAIR10) with STAR (Dobin et al. 2013) in two-pass mode. Alignment parameters included removal of non-canonical intron motifs, a maximum of 20 multimapping loci, a minimum overhang of 8 nt for unannotated and 1 nt for annotated junctions, a maximum mismatch rate of 0.04, an intron size range of 20–100,000 nt, a maximum mate gap of 100,000 nt and a minimum chimeric segment of 12 nt. Alignments were sorted by coordinate and indexed with samtools (Li et al. 2009).

### Intron retention index calculation

Intron coordinates were extracted from the TAIR10 (v59) GTF annotation. For each annotated intron, spliced reads were counted as those containing an N CIGAR operation spanning exactly the annotated 5′ and 3′ splice sites. Unspliced reads were counted independently at the 5′ and 3′ splice sites; a read was scored as unspliced at a given site if it aligned continuously across the exon–intron boundary, covering the splice site and at least one position on each side within a 3-nt window. Read counting was performed with pysam. The intron retention index (IRX) was calculated as IRX = mean(5′ss unspliced, 3′ss unspliced) / [mean(5′ss unspliced, 3′ss unspliced) + spliced reads] × 100, where a value of 100 indicates complete retention and 0 indicates complete splicing. The score was set to NA when total coverage was zero. Retention scores and raw read counts for all introns across all 36 samples were compiled into separate matrices for nuclear and total RNA, with gene annotations retrieved from the TAIR10 GTF.

### Positional analysis of intron retention

The dependence of retention on intron position was analysed separately for total and nuclear RNA, in both genotypes and under all conditions, using the per-intron retention scores together with intron-position annotation. Each intron was assigned an ordinal position counted from the 5′ end (intron number within the transcript) and from the 3′ end (defined as the total number of introns minus the intron number plus one); positions were capped at 10 for display and tabulation. Single-intron genes, whose only intron simultaneously occupies position 1 from both ends, were analysed separately. An intron was included for a given genotype and condition only if its retention index was available for all three replicates and the mean number of informative reads was ≥ 5; the sets of included introns therefore differed between conditions. Genes were additionally stratified by total intron number (1, 2–3, 4–6, 7–10 and > 10 introns).

The relationship between retention and position was quantified by Spearman’s rank correlation computed from each end, and differences in retention among positions were tested with the Kruskal–Wallis test. The localization of the positional effect was examined by (i) comparing retention between consecutive positions with the Mann–Whitney U test (reported as −log₁₀ p), and (ii) computing the cumulative Spearman correlation obtained when progressively more positions were included (the first k positions) from each end. Retention at the first and last introns was compared directly with the Mann–Whitney U test, and the proportion of introns exceeding a retention threshold at each position was compared with position 1 using the chi-square test, with Wilson confidence intervals for the proportions. Changes in retention between conditions (ΔIRX, stress − control) were calculated for the same introns, i.e., introns passing the filter under both compared conditions, and tested at each position with the Wilcoxon signed-rank test with Benjamini–Hochberg correction; 95% confidence intervals of the mean difference were calculated as ±1.96 SE. The genotype × condition interaction was tested by two-way ANOVA on replicate-level mean IRX (n = 3 per group) of introns passing the filter in all four compared groups.

Joint dependence on position from both ends was visualized as two-dimensional matrices of mean retention as a function of 5′ and 3′ position simultaneously. Analyses were implemented in Python using NumPy, pandas, SciPy, matplotlib and seaborn.

### Sequence features and motif analysis of retained introns

To identify intrinsic sequence properties associated with retention, two intron sets were defined from the retention matrices (introns passing a mean-coverage filter of ≥ 5 reads): efficiently spliced introns, with IRX < 10% in all conditions, genotypes and RNA fractions in which they passed the filter (at least three valid values required); and strongly retained introns, with IRX > 90% in at least one condition, genotype or RNA fraction. Introns present in both sets were removed from the spliced set, yielding 24,393 efficiently spliced and 7,648 strongly retained introns; sequences were extracted for 7,610 of the latter. For each intron, the genomic sequence was extracted with ± 50 nt exonic flanks (intron in lower case, flanks in upper case) from the *A. thaliana* genome using the TAIR10 annotations.

The following features were computed for each intron: intron length; intron GC content; the GC content of the 5′ and 3′ exonic flanks and the GC contrast (intron GC minus mean flanking-exon GC, in percentage points); the fraction of AT-rich tetramers (4-mers containing at least three A/T nucleotides), used as a proxy for the UA-richness signal of plant intron definition; mononucleotide composition; the CpG observed/expected ratio; the 5′ splice-site consensus match (fraction of the first six intronic positions matching the *A. thaliana* donor consensus GTAAGT); the 3′ splice-site consensus match (against TGCAGG, spanning the last intronic and first exonic positions); the polypyrimidine-tract strength (pyrimidine, C+T, fraction over the 20 nt upstream of the 3′ splice site); and the splice-site dinucleotide class (GT–AG, GC–AG, AT–AC or other, with non-canonical defined as any class other than GT–AG). Positional nucleotide-composition profiles across the exon–intron boundaries and sequence logos of the splice-site regions were also generated.

Continuous features were compared between the two intron classes with the two-sided Mann–Whitney U test, with effect sizes reported as the rank-biserial correlation coefficient; categorical features were compared with the chi-square test, with Cramér’s V as the effect size. To confirm that compositional differences were not a by-product of intron position, the feature comparisons were repeated within positional strata. Strongly retained introns were additionally classified by condition specificity (constitutive, i.e. IRX > 90% in all conditions in which the intron passed filters; hypoxia-specific; reoxygenation-specific; retained under both stresses but not control; or control-specific) and by genotype specificity (retained in both genotypes, wild-type-only or *ncb-1*-only).

De novo motif discovery was performed with STREME (motif width 4–15 nt; Bailey 2021) from the MEME Suite (Bailey et al. 2015), with a significance threshold of E < 0.05. The analysis was run on intron sequences without exonic flanks, after trimming 20 nt from each intron end to exclude splice-site signals, in both directions (retained versus efficiently spliced introns and the reverse); the control set of efficiently spliced introns was matched to the retained set for length and GC content (standardized mean differences ≤ 0.003). For comparison, the analysis was repeated without end trimming. The frequency of an internal GUAAG donor-like element and of A-rich elements resembling plant polyadenylation signals was compared between retained and matched spliced introns with Fisher’s exact test, and against dinucleotide-shuffled sequences (Supplementary Table S3). Motif significance is reported as the MEME Suite E-value (the expected number of equally enriched motifs arising by chance in a dataset of the same size).

### Statistical analysis

Group comparisons used the two-sided Mann–Whitney U (Wilcoxon rank-sum) test or the Kruskal–Wallis test as appropriate, with effect sizes reported as the rank-biserial correlation (continuous) or Cramér’s V (categorical); correlations were assessed with Spearman’s rank coefficient; and multiple testing was controlled with the Benjamini–Hochberg procedure unless stated otherwise. A threshold of p (or adjusted p) < 0.05 was used throughout unless indicated.

## Results

### Cajal bodies modulate nuclear retention and intracellular distribution of poly(A^+^) RNA

Initially, we examined whether poly(A^+^) RNA nuclear retention occurs during hypoxia in hydroponically grown plants, and whether any differences exist in the spatial organization of the transcriptome at the cellular level between A. thaliana wild-type (WT) and *ncb-1* mutant plants (Fig. 1A-N).

**Fig. 1.**
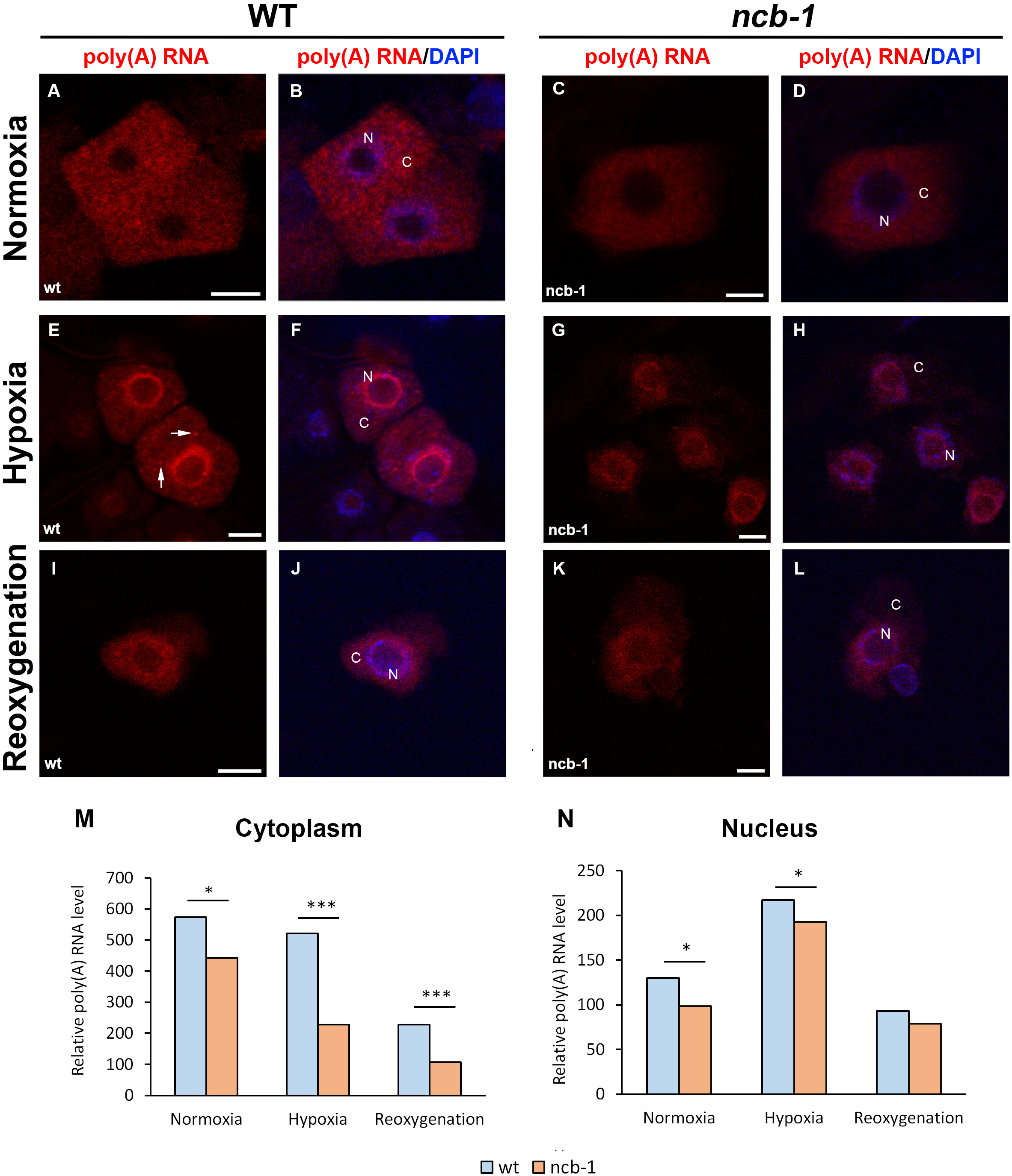
Localization and relative level of poly(A^+^) RNA in root cells of wild-type (WT) *Arabidopsis thaliana* and the *ncb-1* mutant under normoxia, hypoxia, and reoxygenation conditions. Localization of poly(A^+^) RNA in root cells of WT **(A, B, E, F, I, J)** and *ncb-1* (**C, D, G, H, K, L)** under normoxia **(A–D)**, hypoxia **(E–H)** and reoxygenation **(I–L).** Panels A, C, E, G, I, and K show the poly(A^+^) RNA signal (red), whereas panels B, D, F, H, J, and L show the merged images of the poly(A^+^) RNA signal (red) and DAPI-stained DNA (blue). In the images, **N** - nucleus and **C** - cytoplasm, arrows indicate granules in the cytoplasm. **(M)** Relative level of poly(A^+^) RNA in the cytoplasm of WT and *ncb-1* root cells under normoxia, hypoxia, and reoxygenation. **(N)** Relative level of poly(A^+^) RNA in the nucleus of WT and *ncb-1* root cells under the same conditions. Asterisks indicate statistically significant differences between WT and *ncb-1* (*p < 0.05; **p < 0.01; ***p < 0.001). Scale bars: 10 µm.

Under normoxic conditions, the poly(A^+^) RNA signal in the nucleus was homogeneous with few small speckles (Fig. 1A-D). The level of poly(A^+^) RNA in both the cytoplasm and nucleus was higher in WT than in the mutant under normoxia (Fig. 1M, N). During hypoxic stress, poly(A^+^) RNA levels declined in the cytoplasm, while a pronounced accumulation of signal was observed in the nucleus. In WT plants, the poly(A^+^) RNA signal was significantly higher in the cytoplasm and slightly elevated in the nucleus compared to *ncb-1* (Fig. 1E–H, M, N). Notably, poly(A^+^)-containing granules were detected in the cytoplasm of WT plants. Upon reoxygenation, poly(A^+^) RNA levels decreased in both the nucleus and the cytoplasm (Fig. 1I–L). The difference between WT and *ncb-1* was confined to the cytoplasm, while no significant difference was observed in the nucleus (Fig. 1M, N).

Collectively, these results indicate that poly(A^+^) RNA nuclear retention occurs during hypoxia. Furthermore, they suggest the involvement of CBs in this process, as well as in the spatial organization of transcripts in the cell under all examined conditions.

### Cajal bodies regulate transcriptomic responses to hypoxia and recovery

To investigate the contribution of CBs to the hypoxia response, total RNA was isolated from roots of both genotypes under normoxia, after hypoxia and after subsequent reoxygenation. The fold change (FC) was then analyzed between normoxia vs. hypoxia and hypoxia vs. reoxygenation conditions. A total of 4,179 genes showed increased expression in both genotypes during hypoxia (Fig. 2A, Supplementary Table S1). The most highly expressed genes during hypoxia include transcription factors associated with abiotic stress (*HFR1*, *BHLH119*, *MYB27*), proteins involved in sugar metabolism (*SUGTL5*, *SUS3*, *BGLU30*), as well as canonical hypoxia markers. Core hypoxia-responsive genes (Mustroph et al., 2009) were induced more strongly in WT than in *ncb-1* (median log₂FC 1.35 vs. 1.05; higher in WT for 33 of 48 genes; Wilcoxon signed-rank test, P = 0.0015; Supplementary Table S1). Among these genes, our analysis identified several associated with the GO:BP terms related to hypoxia, oxygen level, salicylic acid, and ethylene, indicating a strong response to oxygen deprivation (Supplementary Table Fig. S1A). A significant difference was demonstrated in the number of stress-induced genes between the WT and *ncb-1*. Notably, a substantially higher number of protein-coding genes were upregulated in response to hypoxia exclusively in WT (1,825) than exclusively in *ncb-1* (577), suggesting a direct or indirect role of CBs in transcriptional activity (Fig. 2A; Supplementary Table S1). In addition to shared GO:BP terms, several terms exclusively enriched in WT plants are associated with mRNA processing, splicing, and transesterification reactions, terms absent in the *ncb-1* mutant (Fig. S1B). During hypoxia, 5,192 genes were also downregulated in both WT and *ncb-1* (Fig. 2A, Supplementary Table S1). GO:BP enrichment analysis revealed that downregulated genes were significantly associated with cell wall biogenesis, ribonucleoprotein complex biogenesis, and the mitotic cell cycle (Fig. S1C). Furthermore, GO:CC analysis revealed a strong association of downregulated genes with the cytoskeleton and ribosomes, consistent with reduced translational activity during hypoxia (Fig. S1D).

**Fig. 2.**
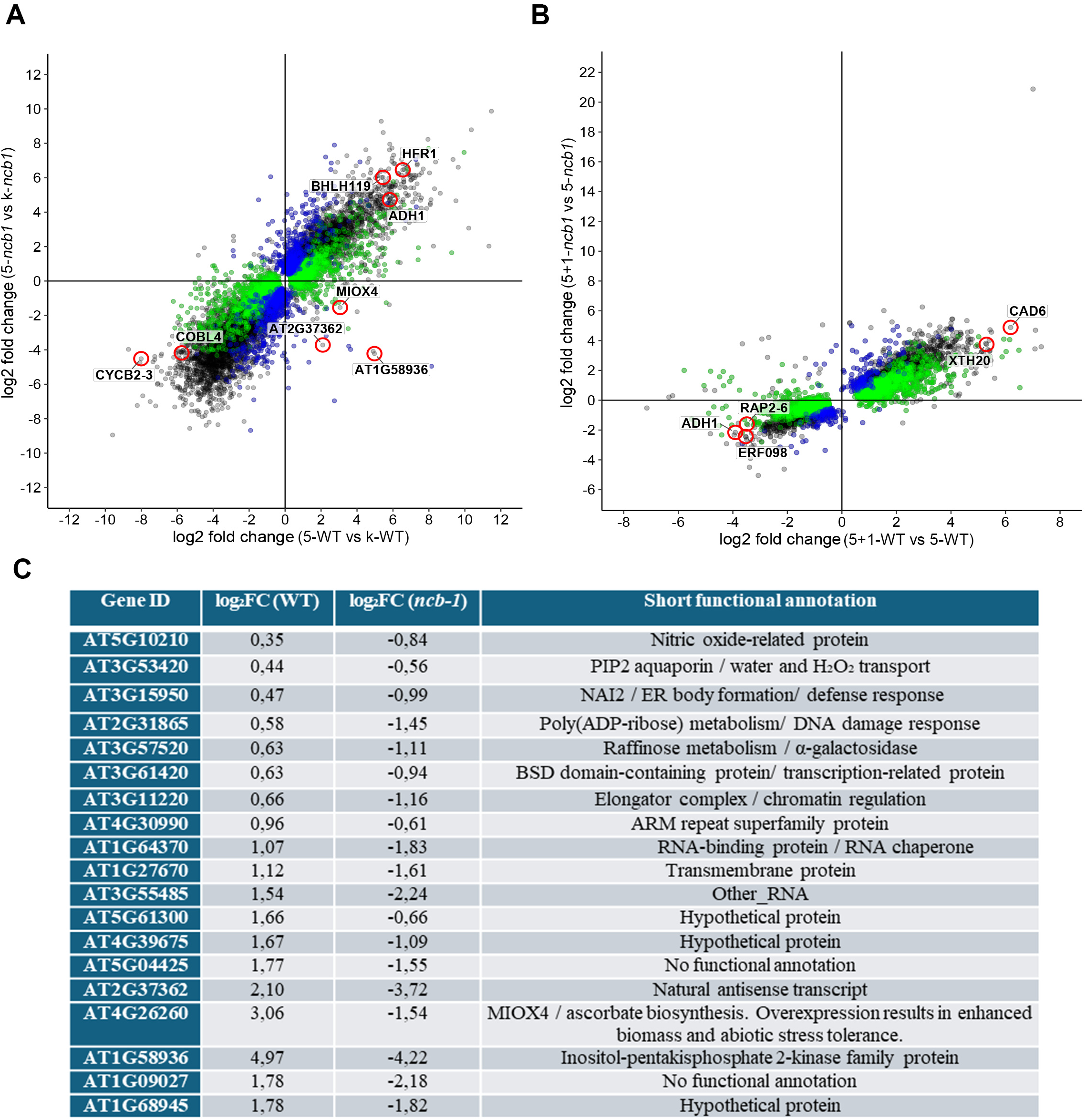
Comparison of differentially expressed genes in roots of WT and the *ncb-1 Arabidopsis thaliana* mutant during hypoxia and reoxygenation. **(A)** Scatter plot of log₂ fold changes after 5 days of hypoxia relative to control conditions. The x-axis represents log₂ fold change values for WT (5-wt vs k-wt), whereas the y-axis represents the *ncb-1* mutant (5-*ncb-1* vs k-*ncb-1*). **(B)** Scatter plot showing changes in gene expression after 1 day of reoxygenation following 5 days of hypoxia compared with 5 days of hypoxia. The x-axis represents log₂ fold change values for WT (5+1-wt vs 5-wt), whereas the y-axis represents *ncb-1* (5+1-ncb vs 5-ncb). **(C)** Summary of selected genes showing divergent expression changes, increase in WT and decrease in *ncb-1* during hypoxia. In (**A**) and (B), green, blue and grey points denote genes differentially expressed (adjusted P < 0.05) only in WT, only in *ncb-1* and in both genotypes, respectively; selected genes are circled in red.

A distinct subset of genes exhibited opposing expression changes between the two genotypes during hypoxia. Given that *ncb-1* is less tolerant to stress than WT (Niedojadło et al., 2016), these genes may represent novel candidate regulators of the hypoxia response. Nineteen genes showed increased expression in WT and simultaneous decreased expression in *ncb-1*. Among those with most significant log₂FC differences were a natural antisense transcript at a locus associated with suberin biosynthesis (AT2G37362), *MIOX4* (AT4G26260), encoding a myo-inositol oxygenase involved in the synthesis of UDP-GlcA, a precursor for cell wall biosynthesis, and of ascorbate, and an inositol-pentakisphosphate 2-kinase family gene linked to phosphate homeostasis (AT1G58936) (Fig. 2C). This suggests a significant role of coilin and CBs in the regulation of transcription, including the response to hypoxic stress.

Transcriptomic changes during reoxygenation relative to hypoxia were subsequently analyzed. Of all genes examined, 912 showed increased expression during reoxygenation, while 597 exhibited reduced expression in both genotypes (Fig. 2B, Supplementary Table S1). In contrast, 1,210 genes were upregulated, while 1,310 were downregulated exclusively in WT. A substantial differences between genotypes was observed, as only 146 genes were upregulated and 107 downregulated exclusively in *ncb-1*, highlighting the significant contribution of CBs to transcriptional regulation during reoxygenation. Among the genes with the greatest expression increases in both genotypes, notable examples include those encoding cyclin B2-3 (*CYCB2-3*, log₂FC = 7.08), cinnamyl alcohol dehydrogenase 6 (CAD6, log₂FC = 6.18), and xyloglucan hydrolase (*XTH20*, log₂FC = 5.30) (Fig. 2B, Supplementary Table S1). GO:BP analysis revealed enrichment of terms primarily related to cell wall biogenesis and metabolism, as well as cell division and fatty acid metabolism (Fig. S1E). Among genes with the greatest upregulation specific to WT, those involved in carbohydrate metabolism (GAE4, log₂FC = 6.52), cell wall synthesis (*IRX9*, log₂FC = 6.49), and small GTPase signaling (*ARAC9*, log₂FC = 5.60) were predominant (Supplementary Table S1). GO:BP analysis indicated strong reactivation of genes that had been downregulated during hypoxia (Fig. S1E); the enriched terms were largely similar in both genotypes; however, WT additionally showed upregulation of genes associated with pyruvate metabolism, ATP, and glycolysis (Fig. S1F). Notably, GO:CC terms for both genotypes were significantly enriched for ribosome-related categories (Fig. S1G-I), although several terms related to mitochondria were additionally identified in genes upregulated exclusively in WT (Fig. S1H).

Among genes showing more than 4-fold downregulation in both genotypes, transcription factors (*WRKY55*, log₂FC = −3.45; *NAC019*, log₂FC = −3.45), an ethylene response factor (*ERF098*, log₂FC = −3.53), and alcohol dehydrogenase (*ADH1*, log₂FC = −3.91) were identified, characteristic of transcriptomic reprogramming following the termination of hypoxia (Fig 2B, Supplementary Table S1). This indicates a reversal of the expression profile during reoxygenation compared to hypoxia: stress-response genes were downregulated. These changes were more pronounced in plants with CBs (e.g., *ADH1*: log₂FC = −3.91 in WT vs. −2.16 in *ncb-1*; Supplementary Table S1), pointing to the involvement of this structure in recovery after stress as well.

The results indicate a role for CBs in shaping transcriptomic profiles during and after hypoxia, revealing markedly greater dynamics of these processes in WT plants. Gene expression analysis showed that hypoxia activates stress-response genes while simultaneously suppressing processes related to cell division and translation. Notably stronger gene activation was observed in the wild type than in the *ncb-1* mutant. Furthermore, the exclusive presence of mRNA processing-related pathways in WT reveals, that CBs play an important role in regulating the molecular response of plants to hypoxia.

### Cajal bodies modulate the composition of the nuclear transcriptome during hypoxia

As demonstrated by our previous studies, CBs influence the nuclear accumulation of total poly(A^+^) RNA and individual transcripts during hypoxia (Niedojadło et al., 2014, 2016). Therefore, RNA-seq was performed on cell nuclei isolated (Isolation of Nuclei Tagged in specific Cell Types; INTACT) from roots under the examined conditions. FC hypoxia/normoxia for both genotypes revealed 614 genes whose nuclear transcript levels increased during stress in both genotypes (Fig. 3A). The first three enriched GO terms were associated with response to decreased oxygen levels, response to oxygen levels, and response to hypoxia (Fig. S2A). This indicates that nuclear mRNA accumulation is linked to increased transcription of hypoxia-responsive genes. Other significantly enriched biological processes are involved in the regulation of gene expression and include chromatin organization, chromatin remodeling, RNA processing, and RNA splicing. These findings were further validated by FISH targeting exonic sequences of highly transcribed genes associated with stress responses and chromatin regulation. Localization of *ADH1* (a hypoxia marker) revealed an increase in mRNA abundance in both the nucleus and cytoplasm under hypoxic conditions (Fig. 3B). However, in *ncb-1* mutants, cytoplasmic mRNA levels during stress were markedly reduced compared to nuclear accumulation, suggesting that despite high transcriptional activity, a larger fraction of this mRNA remains in the nucleus. During reoxygenation, a decrease in ADH1 mRNA levels throughout the cell was observed, consistent with the known biological function of this gene. These results are in agreement with NGS-based nuclear and total transcriptome data (Supplementary Table S1). Another example of a stress-responsive gene associated with chromatin modification is a histone deacetylase gene (*HDA17*) (Fig. 3C), which similarly displayed elevated mRNA levels in both the nucleus and cytoplasm, further supported by NGS data. This suggests that the nuclear transcriptome in both the wild-type and the *ncb-1* reflects a strong plant response to stress, as well as changes in the gene expression profile. A similar stress-dependent redistribution was observed for the BHLH transcript examined by FISH (Fig. 3D).

**Fig. 3.**
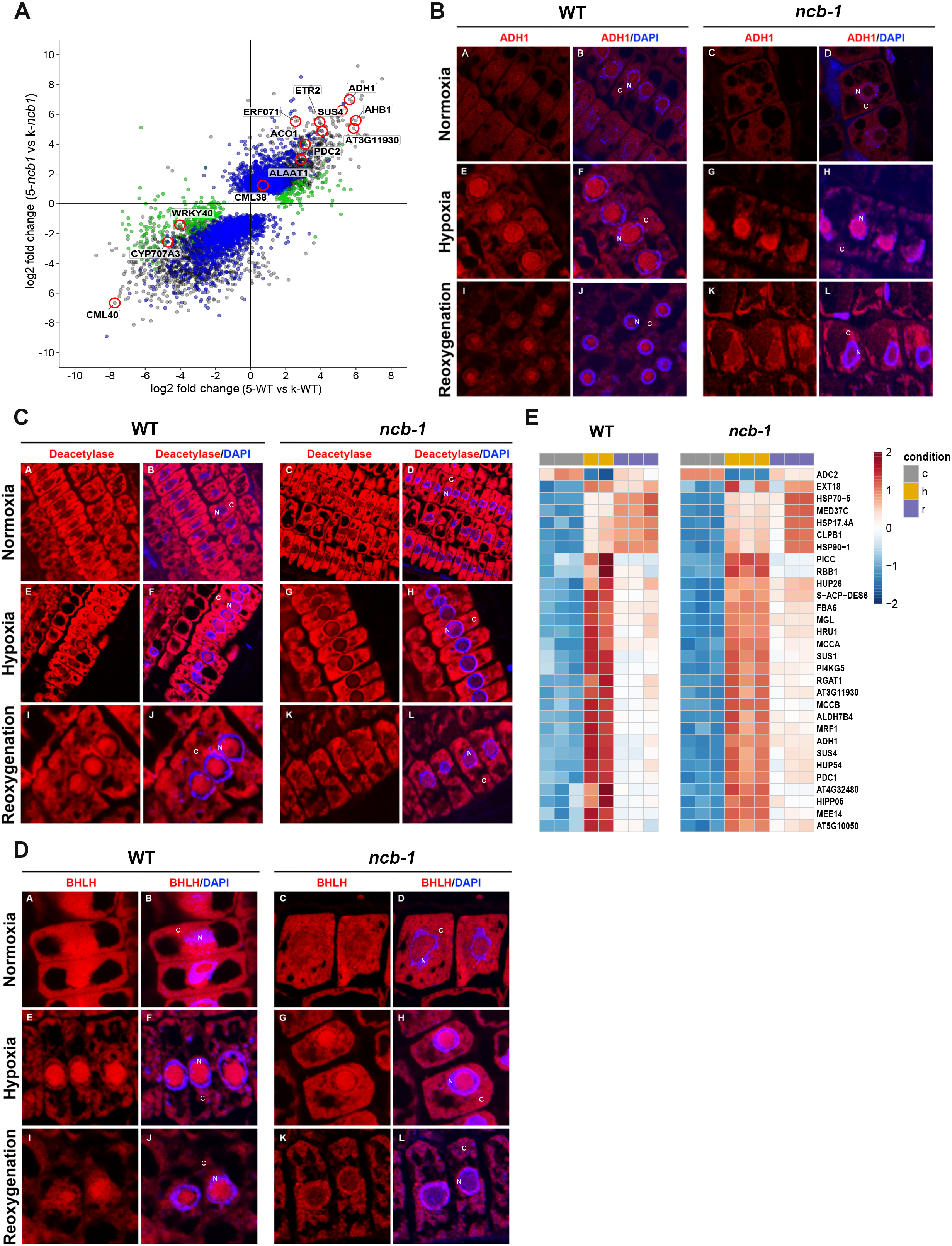
Changes in the expression and localization of selected mRNAs in roots of wild-type (WT) *Arabidopsis thaliana* and the *ncb-1* mutant under normoxia, hypoxia, and reoxygenation conditions. **(A)** Scatter plot of log₂ fold changes in nuclear RNA after 5 days of hypoxia relative to control conditions. The x-axis represents log₂ fold change values for WT (5-wt vs k-wt), whereas the y-axis represents log₂ fold change values for *ncb-1* (5-ncb vs k-ncb). Selected genes are circled in red and labelled; green, blue and grey points denote genes differentially expressed (adjusted P < 0.05) only in WT, only in *ncb-1* and in both genotypes, respectively. **(B–D)** Localization of *ADH1* **(B)**, histone deacetylase *HDA17* **(C)** and *BHLH* **(D)** mRNAs in root cells of WT and *ncb-1* under normoxia, hypoxia, and reoxygenation; within each panel, sub-panels A–L are arranged as in Fig. 1. The mRNA signal is shown in red; in merged images, DNA was stained with DAPI (blue). **N** - nucleus; **C** cytoplasm. **(E)** Heatmaps showing relative expression levels (row z-scores) of selected genes in nuclear RNA of WT and *ncb-1* under control (c), hypoxia (h) and reoxygenation (r) conditions.

The analyses also revealed substantial differences in the nuclear transcriptome depending on the presence of CBs. A total of 119 genes showed statistically significant increases in mRNA levels exclusively in WT, and 1,447 exclusively in *ncb-1* (Fig. 3A, Supplementary Table S1). GO:BP analysis linked genes highly expressed only in WT to stress responses, including hypoxia, and to energy crisis, as evidenced by the following enriched terms: generation of precursor metabolites and energy, pyruvate metabolic process, and isoprenoid metabolic process (Fig. S2B). Stronger expression in *ncb-1* was also observed within the same terms, but frequently for different genes than in WT, including genes associated with hypoxic stress, splicing, and chromatin remodeling (Fig. S2C). This indicates that these are the same categories as those identified for genes whose expression increased in both genotypes.

It was also observed that 568 genes were strongly downregulated during hypoxia in both genotypes. Among the most strongly downregulated genes are those regulating phosphate homeostasis (*AT2G45130*, *SPX3*), a calmodulin-like calcium-binding protein (*AT3G01830*), and tyrosine aminotransferase associated with responses to stress signals namely jasmonates and wounding (*AT2G24850*) (Fig. 3A, Supplementary Table S1). GO analysis primarily revealed terms related to the cell wall, secondary metabolite biosynthetic processes, and glucosinolate metabolic processes (Fig. S2D). Differences were also detected in the sets of downregulated genes between genotypes: 249 in WT and 1,833 in *ncb-1* (Supplementary Table S1). According to GO:BP, in WT these genes are associated with jasmonate signaling and immune responses (Fig. S2E), whereas in *ncb-1* they concern cell wall biogenesis, including the regulation of polysaccharides, pectins, homogalacturonan, and lignin (Fig. S2F).

Genes selected for the heatmap displayed a consistent direction of expression change across both genotypes (WT and *ncb-1*), using for each gene the comparison (H-vs-K or R-vs-K) with the lower padj, then ranked in ascending order of that value regardless of baseline expression level (Fig 3E). Of the 30 genes represented, only a subset (*ADH1*, *PDC1, SUS1*, *SUS4*, *ALDH7B4*, *FBA6*, *HUP26*, *HUP54*, *HRU1*) correspond to established components of the core hypoxia-responsive gene set (Fig. 3E) (e.g., Mustroph et al., 2009). The remaining genes, including several heat-shock/chaperone-related loci (*HSP70-5*, *HSP90-1*, *HSP17.4A*, *CLPB1*, *MED37C*) and genes without a prior hypoxia annotation (*EXT18*, *ADC2*, *S-ACP-DES6*, *MGL*, *PI4KG5*, *RGAT1*, *PICC*, *RBB1*, *MRF1, HIPP05, MEE14, MCCA, MCCB, and three uncharacterized loci*), nonetheless displayed the same expression profile as the canonical hypoxia genes in both genotypes: low under normoxia, induced under stress, and relaxing toward baseline under reoxygenation. At the nuclear transcriptome level, this shared directionality across both categories of genes indicates that the transcriptional response whether narrowly hypoxia-specific or not is preserved in the *ncb-1* mutant. However, the magnitude of the response differed between genotypes: for core hypoxia-responsive genes, induction in the nuclear fraction was stronger in *ncb-1* than in WT (median log₂ FC 3.91 vs. 3.10; lower in WT for 42 of 51 genes; Wilcoxon signed-rank test, P = 2.8 × 10⁻⁷; Supplementary Table S1), in contrast to total RNA, in which it was stronger in WT.

To determine which transcripts may translocate between the nucleus and the total RNA pool under the examined conditions, a further analysis was performed. Comparing normoxia with hypoxia, among transcripts relatively enriched in the nuclear fraction under normoxia and shifted towards the total RNA pool under stress, 100 and 127 transcripts were identified in WT and *ncb-1*, respectively (Fig. 4A, B, Supplementary Table S2). Thirty-seven of these were common to both genotypes. Notably, over 30 are associated with stress responses, including 5 specifically related to hypoxia (Supplementary Table S2). This suggests that nuclear retention may represent a regulatory mechanism that primes plants for potential stress. In turn, 55 transcripts in WT and 42 in *ncb-1* became relatively enriched in the nuclear fraction during hypoxia (Fig. 4A, B, Supplementary Table S2). In this case, only 10 transcripts were shared between the two genotypes, suggesting a genotype-dependent component of this process. These are linked to primary metabolism, including ribosomal proteins, transport, and cellular homeostasis. The main differences between these two gene sets concern the substantially greater number of transcripts involved in cell wall remodeling in WT (Supplementary Table S2). In the subsequent step, transcripts shifting from the nuclear fraction towards the total RNA pool during reoxygenation were identified. Only 10 such transcripts were found in WT and 4 in *ncb-1*, none of which were shared, whereas 13 and 24 transcripts, respectively, showed the opposite shift (4 shared; Fig. 4C, D, Supplementary Table S2). This analysis showed that the accumulation of transcripts in the nucleus may play a role in the regulation of a specific subset of genes under hypoxic conditions. Differences between the WT and *ncb-1* lines also suggest that CBs are involved in nuclear retention.

**Fig. 4.**
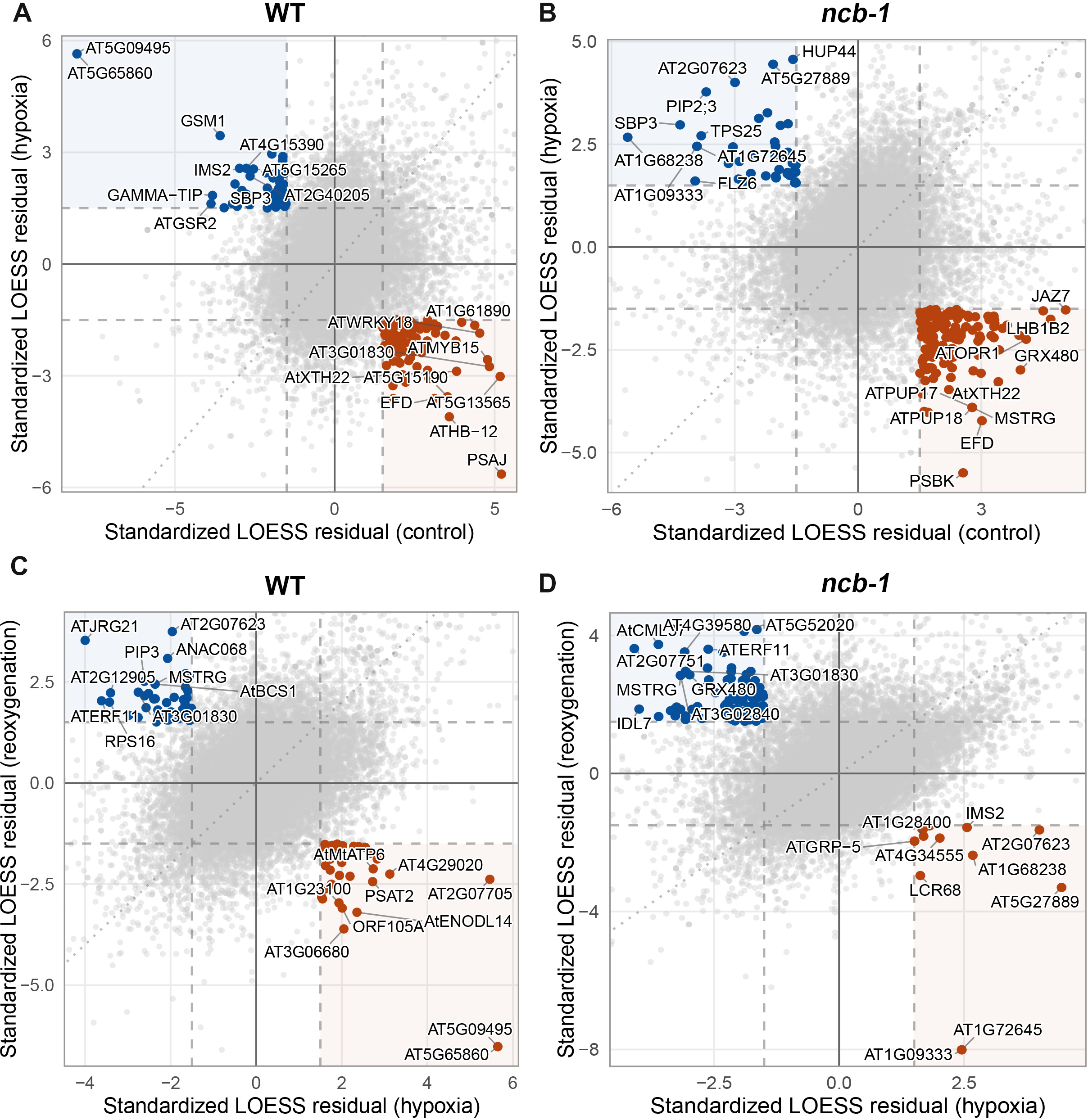
Comparison of standardized LOESS residuals for the analysed genes in roots of wild-type (WT) *Arabidopsis thaliana* and the *ncb-1* mutant under control, hypoxia, and reoxygenation conditions. **(A, B)** Scatter plots showing the relationship between standardized LOESS residuals under control conditions (x-axis) and hypoxia (y-axis), for **WT (A)** and the *ncb-1* mutant (B), respectively. **(C, D)** Scatter plots showing the relationship between standardized LOESS residuals under hypoxia (x-axis) and after reoxygenation (y-axis), for **WT (C)** and the *ncb-1* mutant (D), respectively. Grey points represent all analysed genes, whereas coloured points indicate genes showing opposite standardized LOESS residual values between the compared conditions. **Blue points** represent genes with a negative LOESS residual in the first condition and a positive residual in the second condition, whereas **orange points** represent genes showing the opposite pattern. Standardized residuals were obtained from a LOESS fit (span = 0.5) of log₁₀-transformed nuclear versus total RNA expression; genes were classified as switching when the residual exceeded 1.5 in one condition and was below −1.5 in the other (dashed lines).

### Genome-wide analysis of intron retention

Since nuclear mRNA accumulation is frequently associated with intron-containing isoforms, which may thereby evade degradation, intron retention was examined across the experimental conditions. To determine whether intron retention in *A. thaliana* depends on the ordinal position of the intron within the gene, we computed the intron retention index (IRX) for each intron and analysed its relationship to position counted from both the 5′ and 3′ ends. The analysis was performed on total and nuclear RNA fractions from WT and *ncb-1* plants under normoxic control, hypoxia, and reoxygenation conditions. Single-intron genes were analysed separately, as their sole intron occupies position 1 from both ends simultaneously.

### Terminal and first introns are preferentially retained

Across all conditions, genotypes, and RNA fractions, intron retention showed a significant negative correlation with intron position from both the 5′ end (Spearman ρ = −0.04 to −0.19, p < 10⁻⁶ in all comparisons) and the 3′ end (ρ = −0.09 to −0.32, p < 10⁻⁴²), with the 3′ correlation stronger than the 5′ correlation in every comparison (1.2–3.1-fold, median 1.7-fold; the asymmetry was largest in the nuclear fraction) (Fig. S3A, B). Retention differed significantly between intron positions in all comparisons (Kruskal-Wallis H = 36–4494, p < 10⁻⁴). In total RNA, mean IRX at the last intron (position 1 from the 3′ end) was approximately 20–24%, dropping sharply to 10–14% at the penultimate position (Fig. 5A, B), whereas from the 5′ end the first two introns showed similarly elevated retention (∼14–18%) with a more gradual decline at interior positions (Fig. 5A-D).

**Fig. 5.**
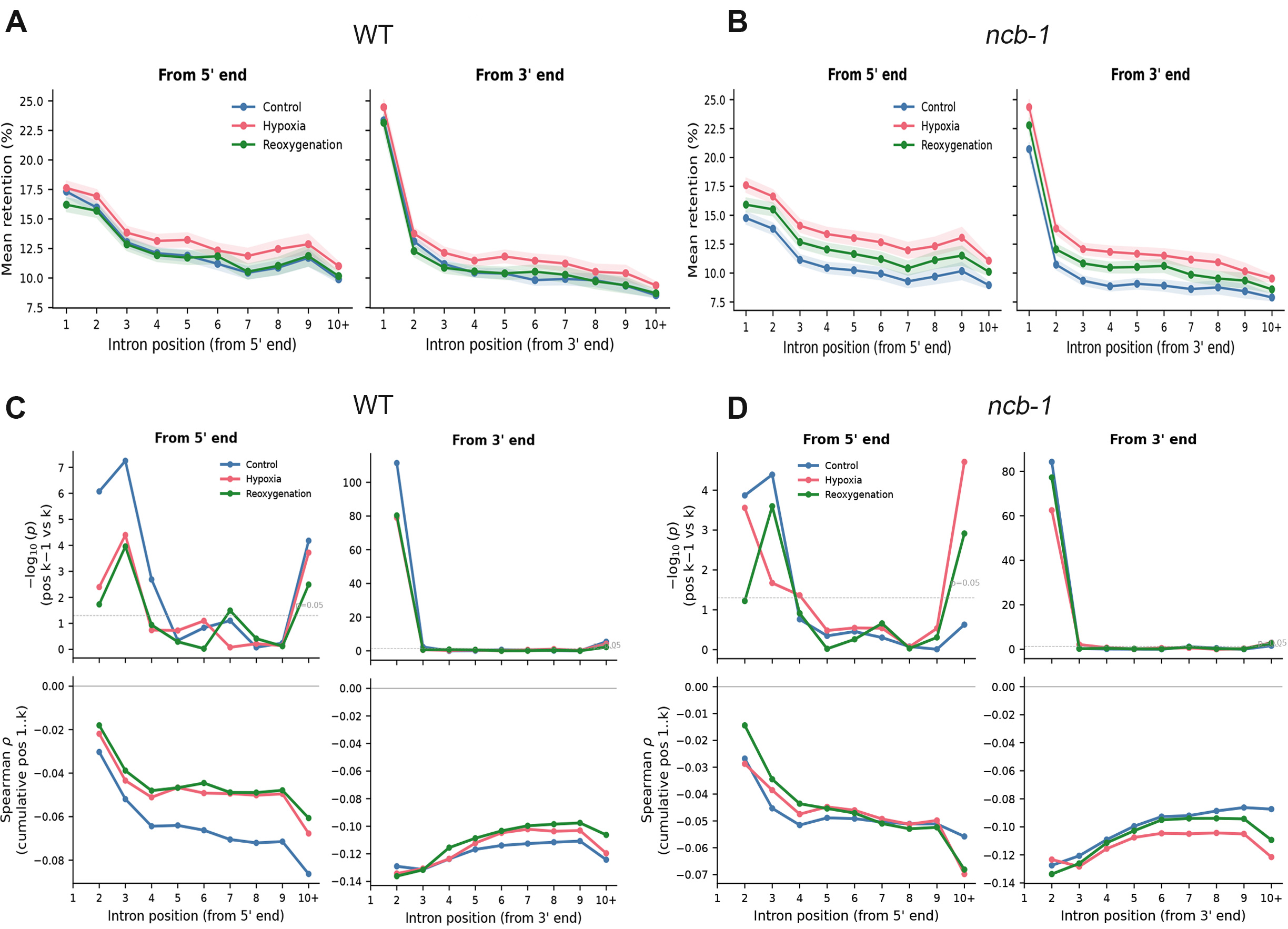
Relationship between intron retention and intron position in total RNA. **(A, B)** Mean intron retention index (IRX) in total RNA according to intron position, counted from the 5′ end and from the 3′ end, for **WT (A)** and the ***ncb-1* mutant (B)**, respectively. Intron positions are shown consecutively from the first to the tenth and subsequent positions (10+). The lines represent values obtained under control conditions (**blue**), during hypoxia (**red**), and after reoxygenation (**green**). Shaded areas represent 95% confidence intervals. **(C, D)** Analysis of the relationship between intron position and intron retention for **WT (C)** and the ***ncb-1* mutant (D)**, respectively. The upper plots show −log₁₀(*p*) values for comparisons between consecutive intron positions (*k*−1 vs *k*), analysed from the 5′ and 3′ ends of the transcript. The horizontal dashed line indicates the statistical significance threshold of *p* = 0.05. The lower plots show the cumulative Spearman rank correlation coefficient (ρ) between intron position and intron retention level, calculated for successive intron positions from the 5′ or 3′ end.

Consecutive-position analysis confirmed that these two effects have fundamentally different profiles. From the 5′ end, the largest drop occurred between the second and third introns (Fig. 5C, D), not between the first and second, indicating that the first two introns share similarly elevated retention before transitioning to baseline at more interior positions. From the 3′ end, the significance of the retention drop was concentrated entirely at the boundary between the last and penultimate introns (Mann-Whitney, −log₁₀(p) = ≈60–110; Fig. 5C, D), with no significant consecutive differences thereafter.

Subsequently, we analysed two-dimensional heat maps showing the mean IRX value as a function of intron position in the nuclear transcriptome (Fig. 6A–F). In WT, IRX markedly decreased during hypoxia and subsequently increased during reoxygenation, suggesting more efficient splicing of the nuclear pre-mRNA pool, or a change in its composition, under stress (Fig. 6A–C). In *ncb-1*, IRX decreased only slightly during hypoxia and increased during reoxygenation (Fig. 6D–F). Under control conditions, the mean IRX per position class ranged from ∼15% to ∼55% in WT, whereas in *ncb-1* it was markedly lower, ranging from ∼2% to ∼30% (Fig. 6A, D). During hypoxia, these values decreased to ∼4–32% in WT and ∼1–29% in *ncb-1*, and after reoxygenation they reached ∼8–45% and ∼5–39%, respectively (Fig. 6). The number of introns with measurable IRX also differed between conditions (Fig. 6); however, because this number depends mainly on the sequencing coverage of individual libraries, it was not interpreted further.

**Fig. 6.**
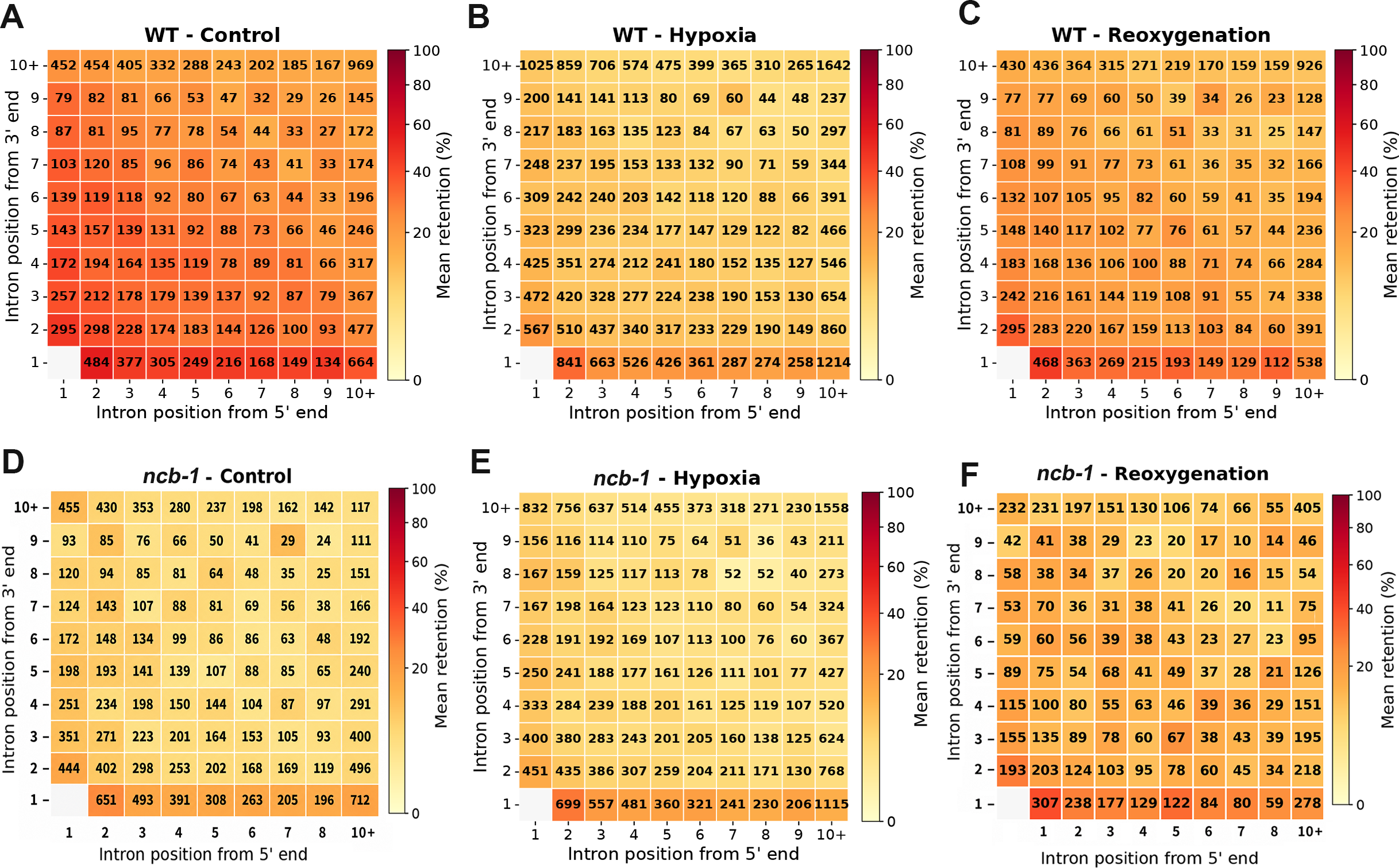
Heatmaps of the mean intron retention index (IRX) in nuclear RNA according to the simultaneous intron position counted from the 5′ end of the transcript (x-axis) and from the 3′ end (y-axis). **(A–C)** Distribution of intron retention in WT under control conditions **(A)**, during hypoxia **(B)**, and after reoxygenation **(C)**. **(D–F)** Corresponding analysis for the *ncb-1* mutant under control conditions **(D)**, during hypoxia **(E)**, and after reoxygenation **(F)**. Intron positions are shown consecutively from the first to the ninth, whereas the **10+** category includes introns located at the tenth or a more distal position from the respective transcript end. The numbers in the heatmap cells indicate the number of introns with measurable IRX (passing the read-coverage filter) in each position class; these numbers depend mainly on the sequencing coverage of individual libraries. The cell corresponding to single-intron genes is empty because these genes were analysed separately.

### Stress modulates intron retention in a compartment dependent manner

To assess the effect of stress independently of differences in sequencing depth between libraries, IRX values were compared for the same introns under control and stress conditions (Fig. 7). In total RNA, hypoxia increased IRX at all intron positions in both genotypes (Δ = +1.7 to +2.4 percentage points in WT and +1.4 to +2.4 in *ncb-1*; Fig. 7A, B), and a smaller increase persisted after reoxygenation (Δ = +1.5 to +2.0 and +0.8 to +1.6, respectively). The magnitude of these changes varied little between intron positions and did not differ between genotypes (genotype × condition interaction for the mean IRX of introns analysed in all four groups: P = 1.0 for hypoxia and P = 0.38 for reoxygenation; two-way ANOVA on replicate-level values, n = 3).

**Fig. 7.**
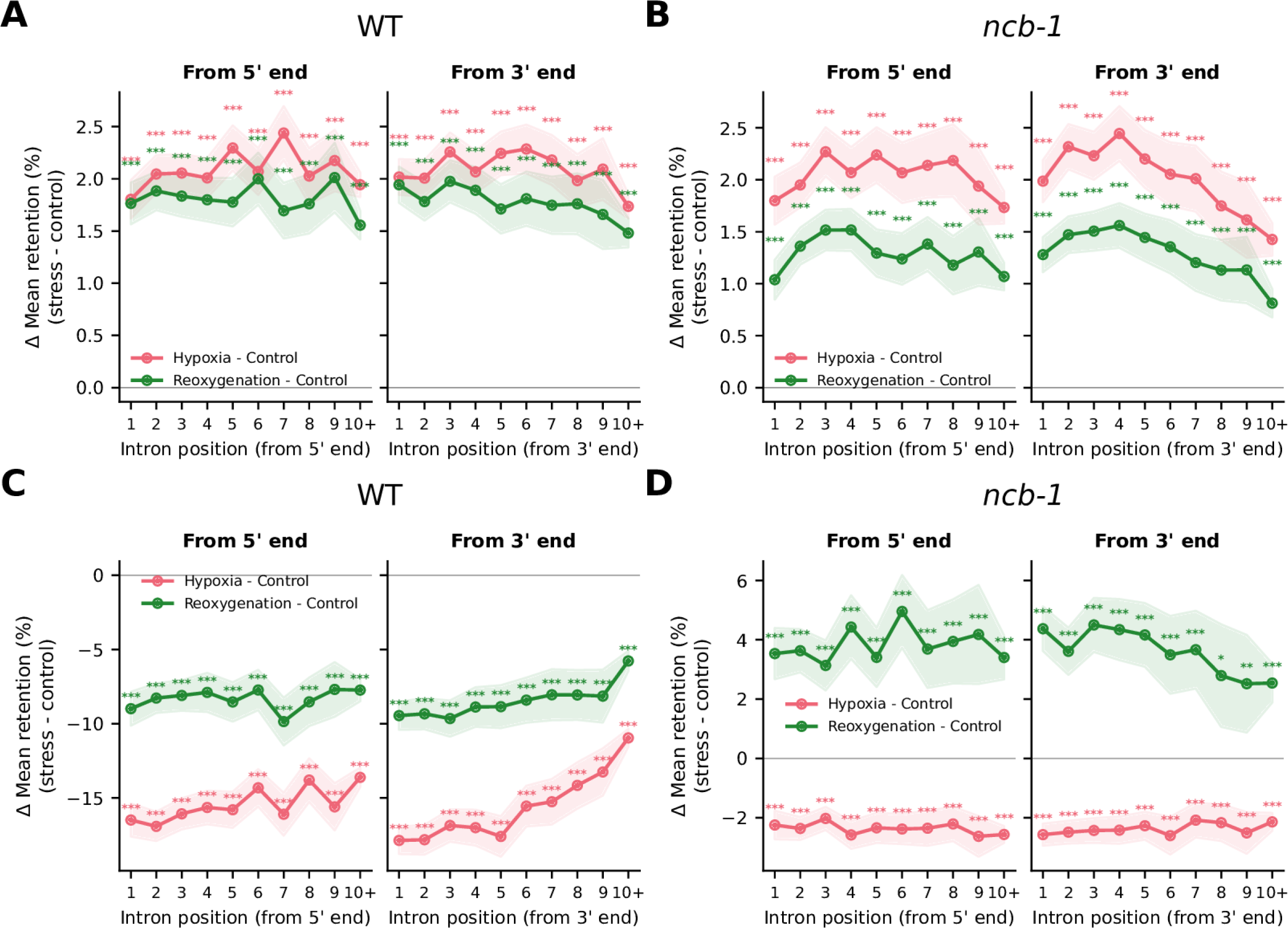
Change in intron retention under stress depending on intron position counted from the 5′ or 3′ end of the transcript, in total (A, B) and nuclear (C, D) RNA of wild-type (WT; A, C) and *ncb-1* (B, D) plants. For each genotype, the difference in the intron retention index (Δ mean retention, stress − control) was calculated for the same introns, i.e., introns passing the read-coverage filter under both compared conditions (multi-intron genes only). Lines show the mean difference for hypoxia − control (red) and reoxygenation − control (green); shaded areas represent 95% confidence intervals of the mean (±1.96 SE). The horizontal line at 0 indicates no difference relative to control conditions. Asterisks indicate the significance of the paired difference at each position (Wilcoxon signed-rank test with Benjamini–Hochberg correction; *P < 0.05, **P < 0.01, ***P < 0.001).

Nuclear RNA revealed a reciprocal pattern; that is, whereas total RNA showed increased retention under stress, nuclear RNA showed decreased retention under the same conditions. Under normoxic and reoxygenation conditions, nuclear IRX was substantially higher than in total RNA (mean ∼24–44% vs ∼16–23% at terminal positions in WT; compare Fig. 6 with Fig. 5), reflecting the expected enrichment for pre-mRNA in the nuclear compartment. In WT, hypoxia reduced nuclear IRX by 11–18 percentage points at all intron positions, and after reoxygenation IRX remained 6–10 points below the control level (Fig. 7C). In *ncb-1*, by contrast, hypoxia reduced nuclear IRX by only 2–3 points, whereas after reoxygenation IRX was 2.5–5 points higher than under control conditions (Fig. 7D). The genotype × condition interaction was significant for both comparisons (P = 0.036 for hypoxia and P = 0.024 for reoxygenation; two-way ANOVA on replicate-level mean IRX) and remained significant after exclusion of one WT control library with an atypical read distribution (P = 0.021 and P = 0.006, respectively). The reversal between total and nuclear RNA is consistent with a model in which hypoxia reduces the transcription rate.

Taken together, these results indicate that the response of intron retention to stress differs between genotypes in nuclear RNA but not in total RNA. In nuclear RNA, the hypoxia-induced decrease in IRX was much stronger in WT than in *ncb-1*, suggesting that CBs contribute to the stress-induced changes in nuclear pre-mRNA processing.

### Gene architecture modulates baseline retention but not positional effects

Stratification by total intron number revealed that baseline IRX decreases with increasing gene intron count, from ∼39–42% in single-intron genes to ∼7–10% in genes with >10 introns in total RNA (Fig. 8A, B), possibly reflecting more efficient spliceosome recruitment in intron-rich, highly expressed genes. Crucially, within each gene-size category, the positional gradient was preserved: the last intron from the 3′ end consistently showed the highest retention, and the first 1–2 introns from the 5′ end were elevated relative to interior positions. In the nuclear fraction, the same hierarchy was observed at higher absolute levels, with single-intron genes reaching ∼62% mean retention in WT under control, compared with ∼39% in *ncb-1* (Fig. S4A, B). Together, these data indicate that the positional pattern of intron retention is independent of gene architecture and is not driven by differences in intron number across the analyzed gene set.

**Fig. 8.**
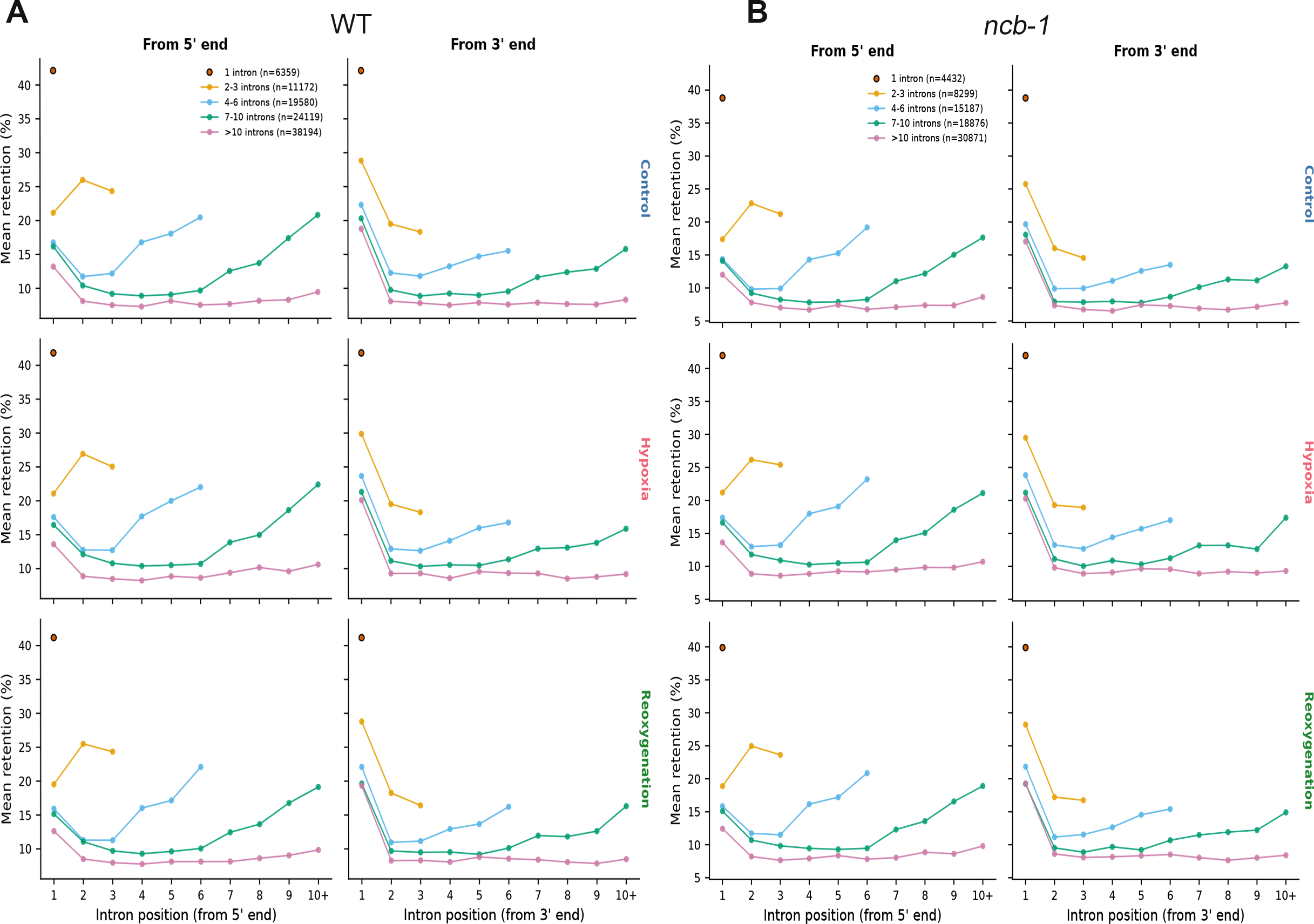
Mean intron retention (IRX) as a function of intron position, calculated separately from the 5′ end and from the 3′ end, in (A) WT and (B) *ncb-1* total RNA. Genes were stratified into five categories by total intron number (1 intron, 2–3, 4–6, 7–10, and >10 introns; n indicates the number of introns analysed per category under control conditions). Rows correspond to condition: control (top), hypoxia (middle), and reoxygenation (bottom). Within each intron-number category, retention is highest for the terminal (3′-most) intron and elevated for the first 1–2 introns from the 5′ end relative to interior positions, while overall retention levels decrease progressively with increasing total intron number.

### Cajal bodies shape the proportions of efficiently spliced and strongly retained introns in nuclear RNA

We next examined how the proportions of efficiently spliced and strongly retained introns changed across conditions. For this purpose, introns were divided into three groups according to IRX: below 10% (efficiently spliced), above 50% and above 90% (the last two groups partly overlapped). The largest differences between conditions were observed for efficiently spliced introns (Fig. 9A, B). In the nuclear RNA of WT plants, introns with IRX < 10% accounted for about 20–49% of the introns detected at a given position under normoxia, but for 61–83% during hypoxia, i.e., an approximately two- to threefold increase. In *ncb-1*, this proportion was already high under normoxia (about 65–83%) and increased only moderately during hypoxia (to about 73–91%). After reoxygenation, the proportion decreased again, remaining above the control level in WT but falling below it in *ncb-1* (Fig. 9A, B). In both genotypes, genes containing efficiently spliced introns were enriched in GO terms related to protein transport, ribonucleoprotein complex and ribosome biogenesis and chromatin organization in nuclear RNA (Fig. S5A, B), and to ribonucleoprotein complex and ribosome biogenesis, RNA processing and cellular respiration in total RNA (Fig. S5C, D). Thus, during hypoxia the nuclear RNA pool of WT plants becomes strongly enriched in efficiently spliced transcripts, whereas this shift is much weaker in the absence of CBs. Consistent with the positional analysis, the last intron (position 1 from the 3′ end) was the least likely to be efficiently spliced in all conditions (Fig. 9A, B).

**Fig. 9.**
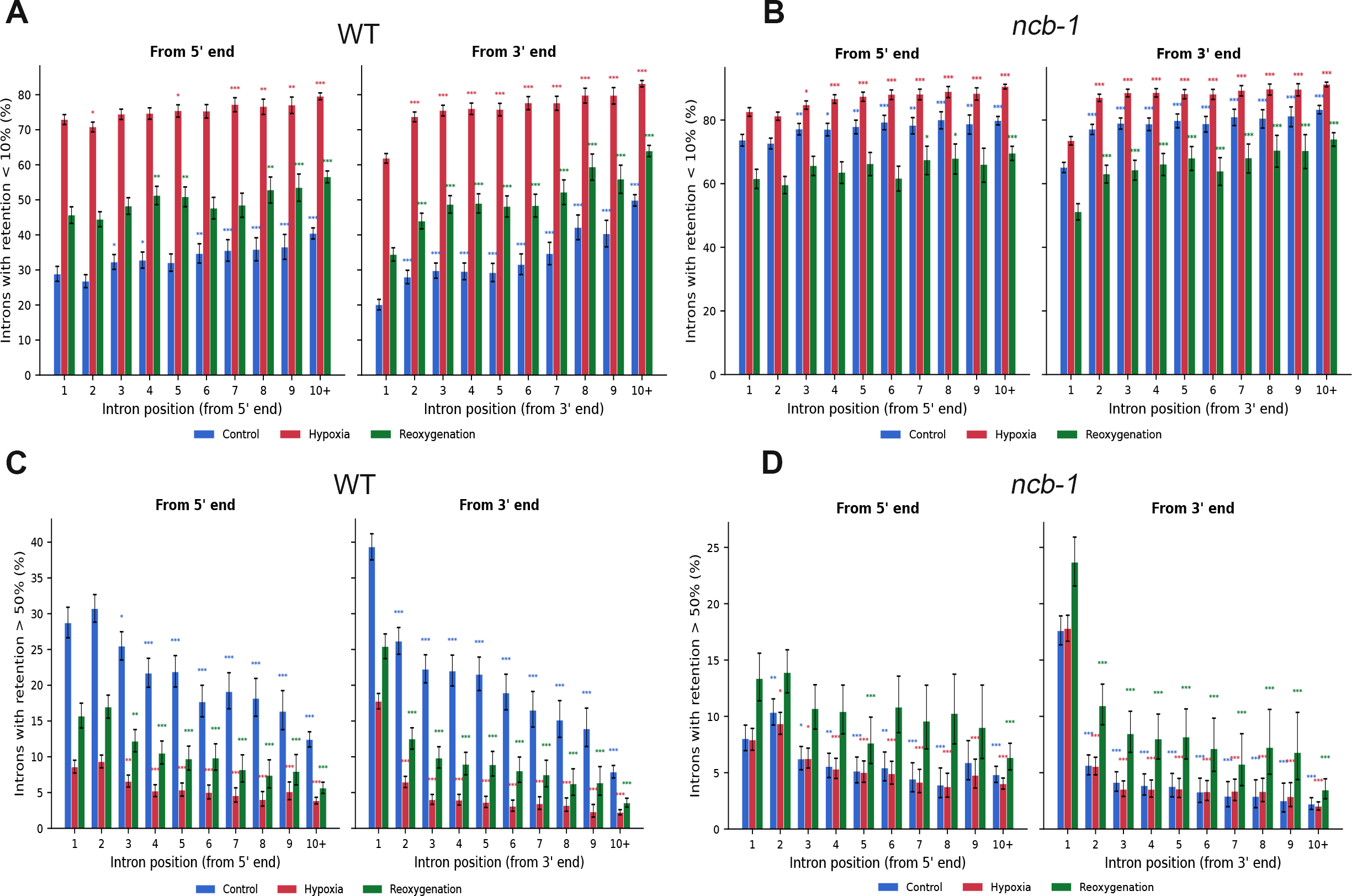
Nuclear retention of introns with low versus high IRX shows opposite dynamics under hypoxia that differ between WT and *ncb-1*. Proportion (%) of introns detected in nuclear RNA that have an intron retention index (IRX) below 10% **(A, B)** or above 50% **(C, D)**, plotted by intron position from the 5′ end (left) and from the 3′ end (right), in **(A, C)** WT and **(B, D)** *ncb-1*. Bars show proportions under control (blue), hypoxia (red), and reoxygenation (green) conditions; error bars represent 95% Wilson confidence intervals. Asterisks indicate statistically significant differences relative to position 1 within the same condition (χ² test; *P < 0.05, **P < 0.01, ***P < 0.001; asterisk colour corresponds to the condition). Only multi-intron genes were included.

The opposite pattern was observed for introns with IRX > 50%. In WT nuclear RNA, their proportion was highest under normoxia (about 8–39%, depending on position), decreased substantially during hypoxia (to about 2–17%) and showed intermediate values after reoxygenation (Fig. 9C). In *ncb-1*, such introns were less frequent under normoxia (about 2–18%), their proportion changed little during hypoxia, and it was highest after reoxygenation (Fig. 9D). Introns with IRX > 90% showed the same strong enrichment at the last intron in both RNA fractions (Fig. S6A–D). In both genotypes, genes containing introns with IRX > 50% in nuclear RNA were enriched in GO terms related to abiotic stress, including responses to water deprivation, decreased oxygen levels and hypoxia (Fig. S7A, B), whereas those identified in total RNA were associated mainly with protein transport, ribonucleoprotein complex biogenesis and DNA repair (Fig. S7C, D). This suggests that, under normoxia, the nuclear RNA of WT plants contains a pool of incompletely spliced transcripts of stress-responsive genes, which is largely depleted during hypoxia. At all conditions, strongly retained introns were more frequent at the last than at the first intron from the 5′ end (Fig. 9C, D).

Taken together, these results show that hypoxia shifts the nuclear pool of intron-containing transcripts in WT plants from incompletely to efficiently spliced forms, and that this shift is strongly attenuated in *ncb-1*, consistent with the changes in mean nuclear IRX (Fig. 7C, D). Because these analyses are based on the proportions of introns detected within the nuclear fraction, they do not by themselves show how transcripts are partitioned between the nucleus and the cytoplasm. Together with the nuclear accumulation of poly(A^+^) RNA during hypoxia (Fig. 1), however, they suggest that largely spliced transcripts are held in the nucleus during stress, whereas incompletely spliced transcripts of stress-responsive genes are present in the nucleus mainly under normoxia. Differences between WT and *ncb-1* were observed in nuclear but not in total RNA (Fig. 7A–D), suggesting that CBs affect intron processing mainly at the nuclear level.

### Most retained introns are retained constitutively

To identify the intrinsic properties associated with intron retention, we compared 24,393 efficiently spliced introns (IRX < 10% in all conditions and genotypes) with 7,648 strongly retained introns (IRX > 90% in at least one condition) (Supplementary Table S3). Of the introns classified, 74.6% were retained under all conditions in which they passed the quality filters, indicating that retention is largely intrinsic to the intron rather than a consequence of stress-induced splicing failure. Stress-specific retention was a minority (6.9% under hypoxia only, 4.2% after reoxygenation only, 4.2% under both). A further 426 introns (5.6%) were retained under control conditions but spliced under stress; these were more often WT-specific (214 WT-only versus 117 *ncb-1*-only), suggesting a reserve of pre-mRNAs held unspliced but readily activatable. Most retained introns were shared between genotypes (83.7%), while 8.1% were confined to *ncb-1* and a similar proportion (8.2%) to WT; the *ncb-1*-specific introns are the strongest candidates for introns whose splicing depends on Cajal body function (Supplementary Table S3).

### Retained introns are compositionally exon-like

The most discriminative feature was not a property of the intron in isolation but its compositional contrast with the flanking exons, which gave the largest effect size of any feature tested (rank-biserial r = 0.38). Efficiently spliced introns were 11.2 percentage points lower in GC content than their flanking exons, whereas retained introns differed by only 7.0 pp (Fig. 10A, Supplementary Table S3). Retained introns thus resemble the surrounding exonic sequence more closely, weakening the UA-rich compositional signal on which intron definition depends in plants (Fig. S8C). The individual composition measures were facets of the same shift: retained introns were more GC-rich (median 34.7% versus 32.7%) (Fig. S8B), poorer in T (38.5% versus 41.4**%)** (Fig. S8D–G) and contained fewer AT-rich tetramers (0.577 versus 0.620) (Fig. S8H). Retained introns were also longer (median 133 versus 103 nt) (Fig. 10B, Fig. S8I).

**Fig. 10.**
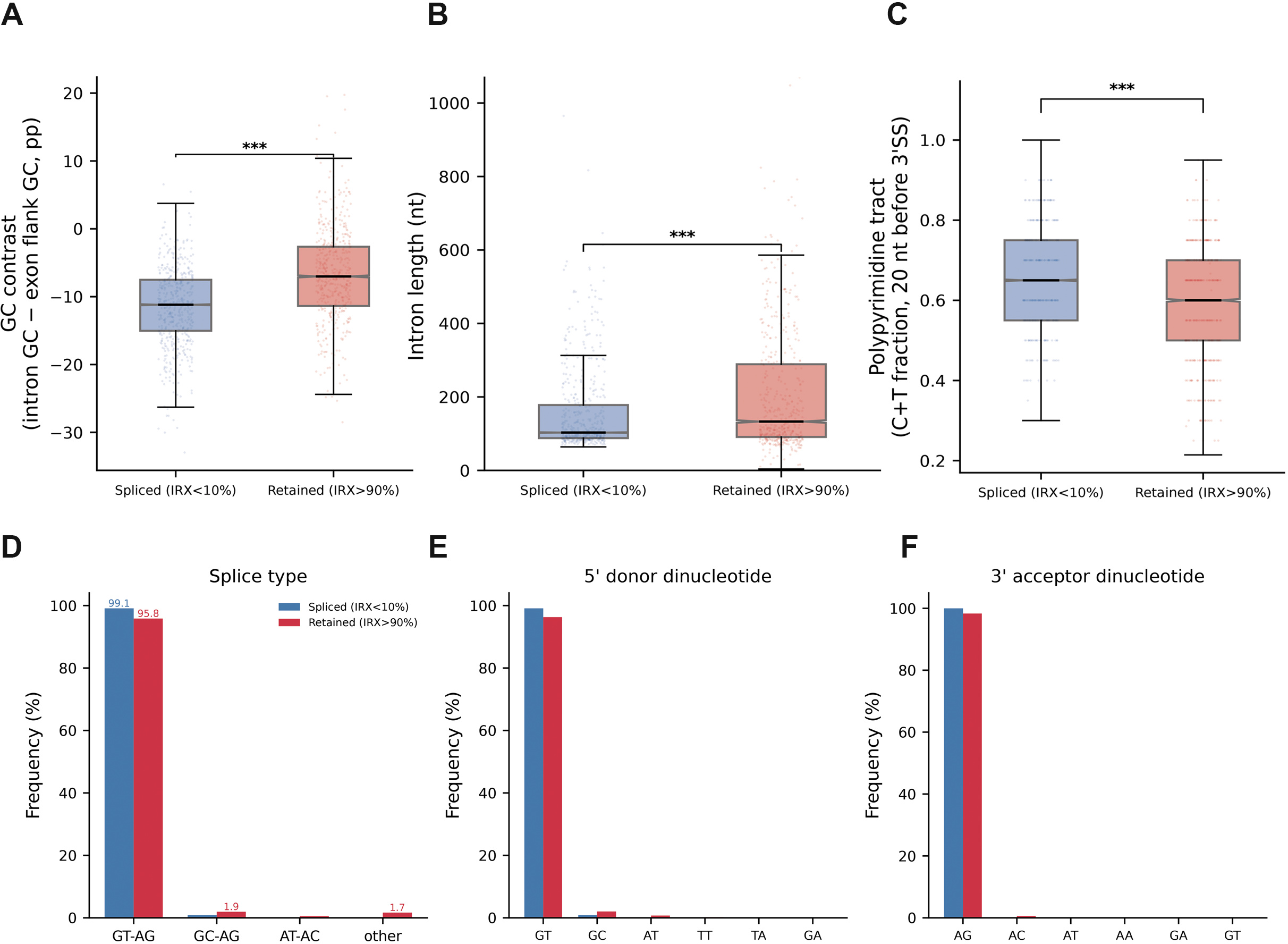
Sequence and structural features of efficiently spliced and strongly retained introns in *Arabidopsis thaliana* roots. **(A)** GC contrast between the intron and its flanking exons, expressed as the difference in GC content in percentage points. **(B)** Distribution of intron length (nt) in efficiently spliced and strongly retained introns. Boxplots show the median and interquartile range, with individual introns shown as points. **(C)** Distribution of polypyrimidine tract strength, expressed as the fraction of C and T nucleotides within the 20-nt region upstream of the 3′ splice site (3′SS). **(D)** Frequency of canonical and non-canonical splice-junction types in efficiently spliced and retained introns. **(E)** Distribution of 5′ donor dinucleotides in the two intron groups. **(F)** Distribution of 3′ acceptor dinucleotides in the two intron groups. Blue represents efficiently spliced introns and red represents strongly retained introns. Asterisks in (A–C) indicate statistically significant differences between the two groups (***p < 0.001).

None of this was a by-product of intron position: the same contrast was observed within every positional class (first, second, internal, last and single-intron genes; Supplementary Table S3, Fig. S9). Length was the exception. Among first introns and single-intron genes the relationship was reversed, with retained introns shorter than spliced ones (152 versus 184 nt and 178 versus 296 nt), consistent with intronic splicing enhancers whose cumulative effect increases with length (Supplementary Table S3).

### Splice signals are weaker in retained introns

Retained introns had weaker polypyrimidine tracts (median pyrimidine fraction 0.60 versus 0.65, r = 0.23; Fig. 10C, Fig. S10A, Supplementary Table S3) and were more than four times as likely to carry terminal dinucleotides other than GT–AG (4.2% versus 0.9%), including the minor GC–AG class (1.9%) and non-canonical combinations (1.7%) (Fig. 10D–F; Supplementary Table S3). The 5′ splice site was also weaker, although the difference was distributional rather than central: median consensus scores were identical (0.67) while the means differed (0.639 versus 0.709, r = 0.21), reflecting a longer tail of poorly matching donor sites among retained introns (Supplementary Table S3, Fig. S10C). Efficiently spliced introns also showed a sharper drop in GC content on entering the intron at both splice sites (Supplementary Table S3; Fig. S10D, E).

### *De novo* motif discovery identifies A-rich elements

Motif discovery was performed with STREME (Bailey 2021) on intron sequences alone, after excluding the flanking exons, trimming 20 nt from each intron end to remove constitutive splice signals, and matching the control set to the retained set for length and GC content (standardised mean differences ≤ 0.003) (Supplementary Table S3). Four motifs were significantly enriched in retained introns (E < 0.05) (Supplementary Table S3) and three in the opposite direction (Supplementary Table S3). The two strongest motifs were A-rich: AAAAAAAAACA (STREME E-value = 7.1 × 10^−3^) and the purine-rich AAARAASARAAN (E = 9.6 × 10^−3^). Their purine character is consistent with binding sites for SR proteins, which could recruit exon-definition factors to intronic positions, while their A-richness resembles the near-upstream element of the plant polyadenylation signal, an A-rich region of 6–10 nucleotides located 10–30 nt upstream of the cleavage site (Loke et al., 2005). In a targeted test, the A-rich octamer AAVAAAAR occurred in 10.5% of retained introns versus 6.3% of length- and GC-matched spliced introns (Supplementary Table S3), 2.2-fold more often than predicted from the dinucleotide composition of the same sequences. As intronic poly(A^+^) signals cause premature cleavage and polyadenylation, they are a plausible contributor to the retention observed here. The remaining motifs were a single CG-containing element (CGAMGA), consistent with the elevated CpG observed/expected ratio in retained introns (Supplementary Table S3; Fig. S10B) and a pyrimidine-rich CU repeat (UCUCUCUUUU) (Supplementary Table S3). Alternating CU tracts of this kind are the preferred binding site of polypyrimidine tract binding protein (PTB), whose specificity is distinct from that of U2AF65 and which can compete with U2AF65 for the polypyrimidine tract, thereby blocking 3′ splice site recognition (Singh et al., 1995; Saulière et al., 2006). *Arabidopsis* encodes three PTB homologues, two of which regulate intron retention transcriptome-wide (Rühl et al., 2012; Simpson et al., 2014), making such competition a plausible mechanism in this system. Motifs enriched in efficiently spliced introns (UAGU, UGCUG, GCAUG; E = 2.2–3.3 × 10^−2^) all contained U and A (Supplementary Table S3), consistent with the role of UA-rich elements as intronic splicing enhancers in plants (Simpson et al. 2004).

Without end-trimming, the most significant motif enriched in retained introns was CAGGUAAG (E = 1.8 × 10^−4^) (Supplementary Table S3), resembling a canonical exon–intron junction. As these sequences contained no exonic flanks, the motif necessarily lies within the intron, pointing to cryptic or alternative 5′ splice sites: an internal GUAAG was present in 12.0% of retained versus 8.7% of spliced introns (odds ratio 1.44, p = 6.4 × 10^−10^) (Supplementary Table S3). Retained introns therefore differ not only in composition and splice-signal strength, but also in harbouring competing donor sites.

### Cajal bodies modulate snRNA and snoRNA expression and promote U2 snRNP assembly in response to hypoxic stress

To describe the potential effect of CB on splicing, we analysed the expression levels of sn- and snoRNAs (Fig. 11A-F) and the assembly of U2 snRNA with U2B″ protein in wild-type and *ncb-1* mutant (Fig. 11G). First, we examined snRNA expression levels under normoxia, hypoxia, and reoxygenation. Overall snRNA expression profiles did not change markedly under hypoxia in either genotype (Fig. 11A-C). Under normoxia, some snRNA genes were differentially expressed between genotypes, suggesting that their expression partly depends on CB status (Fig. 11A). In contrast, under hypoxia, the same subset of snRNA genes is most highly activated in both genotypes, suggesting the existence of stress-responsive snRNAs whose expression is largely unaffected by CBs (Fig. 11B). Of the 8 snRNAs more highly expressed in wild-type than in *ncb-1*, as many as 5 belong to the U1 snRNA class (Fig. 11A, F). Components of the U1 snRNP are known to modulate stress-responsive alternative splicing in *Arabidopsis* (de Francisco Amorim et al., 2018). snoRNA expression was significantly higher in WT than *ncb-1* under normoxic and reoxygenation conditions (Fig. 11E). CB-associated factors participate in snoRNP assembly and in snoRNA- and scaRNA-guided RNA modification (Lermontova et al., 2007; Kannan et al., 2008; Vitali and Kiss 2019). Since CBs are considered not only as structures that regulate snRNA transcription but also as sites of their interaction with proteins, we analysed the function of CBs in the assembly of the U2 snRNA with the U2B″ protein. Both genotypes expressed U2B″–GFP: the *ncb-1* mutant was originally isolated in a U2B″–GFP background (Collier et al., 2006), and the parental U2B″–GFP line was used as the WT control. RNA immunoprecipitation (RIP) of U2B″–GFP was performed with anti-GFP nanobody beads (GFP-Trap), and the amount of U2 snRNA associated with U2B″–GFP was assessed by RT-qPCR in WT and *ncb-1* plants. Western blot analysis confirmed the immunoprecipitation of U2B″–GFP in both genotypes, although the signal in the eluate was weaker in *ncb-1* (Fig. S11). The percentage of input U2.2 and U2.3 snRNA associated with U2B″–GFP was significantly higher in WT than in *ncb-1*, by approximately 2.3-fold and 6-fold, respectively (Fig. 11G).

**Fig. 11.**
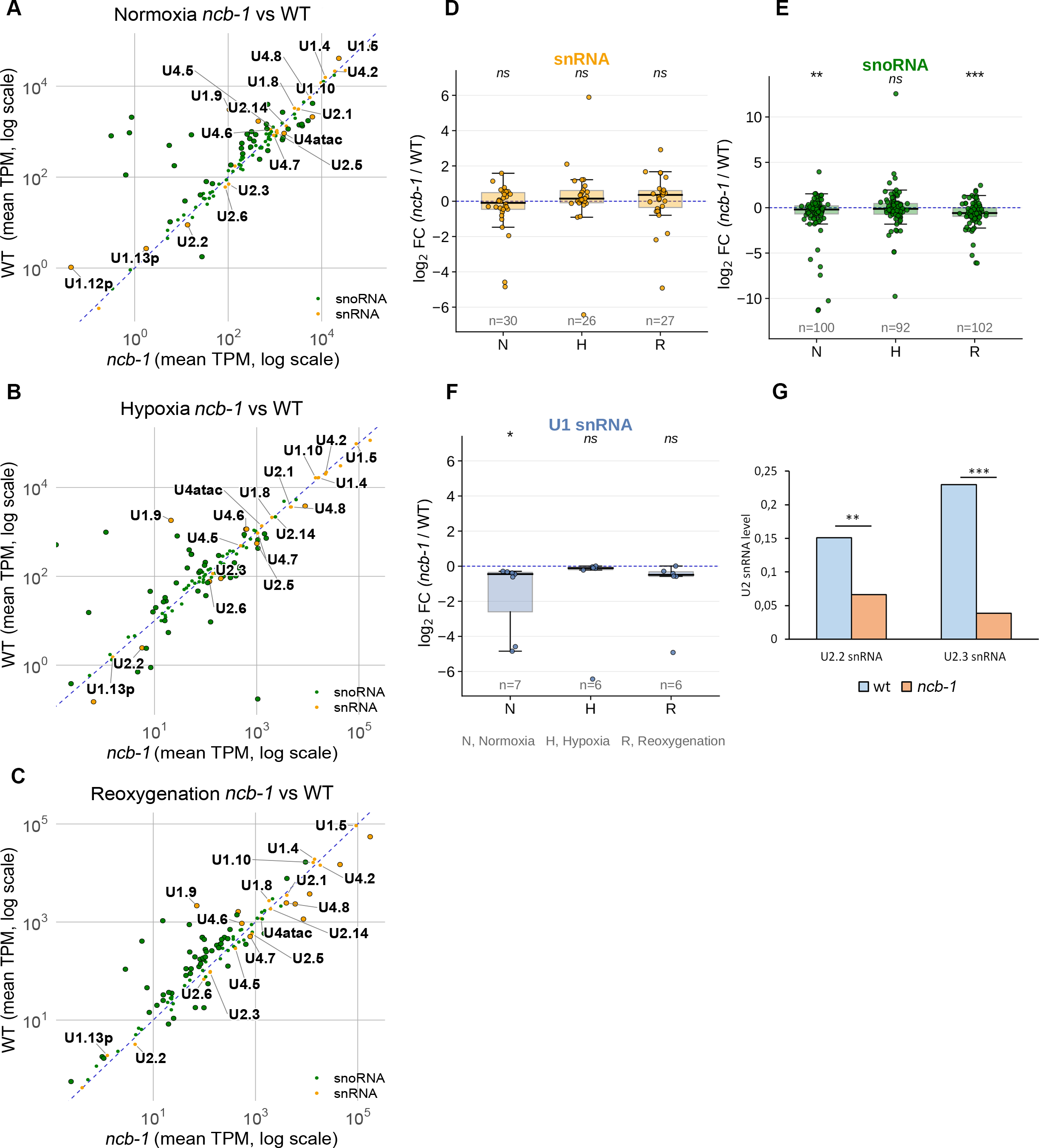
Comparison of snRNA and snoRNA levels and the association of U2 snRNA with U2B″ in the roots of WT *Arabidopsis thaliana* and the *ncb-1* mutant under normoxic, hypoxic, and reoxygenation conditions. **(A–C)** Scatter plots showing the levels of individual snRNAs and snoRNAs, expressed as TPM (transcripts per million; logarithmic scale), in the *ncb-1* mutant (x-axis) and WT (y-axis) under **normoxia (A)**, **hypoxia (B)**, and after **reoxygenation (C)**, respectively. snRNAs are shown in orange, whereas snoRNAs are shown in green. Distribution of log₂ fold change values [log₂FC (*ncb-1*/WT)] for snRNAs **(D)**, snoRNAs **(E)** and U1 snRNAs **(F)** under normoxia **(N**), hypoxia **(H)**, and reoxygenation **(R)**. Positive values indicate higher levels in *ncb-1*, whereas negative values indicate higher levels in WT. The number of analysed RNAs (*n*) is indicated below each group. **(G)** Comparison of the levels of U2 snRNA variants (U2.2 and U2.3) associated with U2B″–GFP between WT and *ncb-1*. Bars represent the levels of the respective U2 snRNA variants in both genotypes. Asterisks indicate statistically significant differences between WT and *ncb-1*, whereas ns indicates no statistically significant difference.

Given these findings, the results suggest that CBs influence snRNA gene activity, including higher U1 snRNA expression in WT. CBs also appear to promote the association of U2 snRNA with U2B″. Furthermore, our results indicate that the *ncb-1* mutant may exhibit disruptions in U2 snRNP as well as sn- and snoRNA expression, which may contribute to disturbances in pre-mRNA splicing.

## Discussion

### The role of Cajal bodies in the hypoxic stress response and transcriptome regulation

Cajal bodies (CBs) have long been regarded as nuclear structures involved primarily in the biogenesis and maturation of snRNPs and snoRNPs (Machyna et al., 2013; Strzelecka et al., 2010; Love et al., 2017). However, a growing body of evidence points to roles extending beyond small nuclear RNP metabolism, including involvement in the response to abiotic stress, nuclear retention of transcripts, regulation of splicing and, more recently, mRNA metabolism (Niedojadło et al., 2016; Górka et al., 2022; Rudzka et al., 2022; Taliansky et al., 2023). The results of the present study provide new, comprehensive evidence for the multifunctional involvement of CBs in the regulation of gene expression during hypoxia and reoxygenation in *Arabidopsis thaliana*.

Transcriptome analysis revealed that hypoxia leads to extensive reprogramming of gene expression in both genotypes analyzed. In total, 4,160 genes were upregulated under oxygen deprivation in both WTplants and the *ncb-1* mutant. At the same time, the number of genes induced exclusively in WT was more than three times higher than in *ncb-1* (1,696 vs. 516 genes), indicating a direct or indirect contribution of CBs to transcriptional activation during stress. The *ncb-1* mutant carries a G→A point substitution in the At1g13030 gene encoding AtCOILIN, which disrupts correct processing of its transcript and leads to the loss of CBs (Collier et al., 2006). Coilin itself may also affect the expression profile, which may be related to the fact that up to 70% of coilin can be dispersed throughout the nucleoplasm in animal model (Lam et al., 2002). Moreover, AtCOILIN has been identified among chromatin-associated proteins in plants (Rayapuram et al., 2021). Abulfaraj et al. (2022) showed that global transcriptome analysis of Atcoilin mutants subjected to biotic stress revealed 667 differentially expressed genes (DEGs) between the mutant and WT under control conditions, 503 of which were upregulated in the mutant. The authors further demonstrated a more altered response to pathogens, indicating that Atcoilin is involved in both transcriptional and post-transcriptional regulation of the defense response. Taken together with the markedly weaker transcriptomic response of *ncb-1* during hypoxia observed here, these results point to a broader, stress-type-dependent involvement of CBs and Atcoilin in modulating gene expression, and suggest that Atcoilin/coilin may have a conserved function in supporting transcriptional activity under both abiotic and biotic stress.

Among the genes induced in WT but repressed in *ncb-1* were AT2G37362, a natural antisense transcript at a locus associated with suberin biosynthesis; *MIOX4* (AT4G26260), encoding a myo-inositol oxygenase involved in the synthesis of UDP-GlcA, a precursor of cell wall components, and of ascorbate; and AT1G58936, an inositol-pentakisphosphate 2-kinase family gene linked to phosphate homeostasis. Suberin is a hydrophobic polymer deposited, among other sites, in root cell walls, and may play an important role in adjusting the barrier properties of tissues to adverse environmental conditions, e.g. in response to soil flooding, where it acts as a barrier limiting the transport of gases and ions (Franke and Schreiber 2007). The stronger activation of these genes exclusively in WT indicates that CBs may be involved in the full activation of specific adaptive gene expression programs during oxygen deprivation. This interpretation is further supported by GO:BP analysis, which revealed enrichment of categories related to RNA processing, splicing and transesterification reactions among genes induced during hypoxia specifically in WT. Furthermore, we suggest that the absence of CBs in the *ncb-1* mutant limits both transcription and transcript maturation, resulting in reduced transcriptomic plasticity during stress.

Our results also point to a novel role for CBs in processes occurring during recovery from hypoxic stress. This observation is consistent with earlier data showing an increased number of CBs in cells with high proliferative activity (Machyna et al., 2013). During reoxygenation, the number of genes induced exclusively in WT was 1,210, more than eight times higher than in *ncb-1* (146), and included genes involved in cell wall biosynthesis, such as *GAE4* (log₂FC = 6.52) and *IRX9* (log₂FC = 6.49). Genes most strongly induced in both genotypes included *CYCB2-3* (log₂FC = 7.08 in WT), encoding cyclin B2-3, *CAD6* (6.18), encoding cinnamyl alcohol dehydrogenase, and *XTH20* (5.30), encoding a xyloglucan endotransglucosylase/hydrolase, although their induction was weaker in *ncb-1* (3.44, 4.89 and 3.75, respectively; Supplementary Table S1). The strong gene activation during reoxygenation in WT suggests that the return to normoxia involves the activation of programs responsible for restoring cellular activity, reorganizing the cell wall and resuming growth, which may not proceed correctly in the mutant lacking CBs.

Notably, the genotype-dependent differences were reversed in the nuclear fraction. Whereas in total RNA more genes changed expression during hypoxia exclusively in WT than exclusively in *ncb-1*, in nuclear RNA far more genes changed exclusively in *ncb-1* (Fig. 3A; Supplementary Table S1), and core hypoxia-responsive genes were induced more strongly in the nuclear fraction of *ncb-1* than in that of WT. Similarly, when the same introns were compared between conditions, hypoxia reduced nuclear IRX by 11–18 percentage points in WT but by only 2–3 points in *ncb-1*, and after reoxygenation nuclear IRX remained below the control level in WT but exceeded it in *ncb-1* (Fig. 7C, D), whereas no such genotype difference was found in total RNA (Fig. 7A, B). However, fully spliced mRNAs of core hypoxia-responsive genes were less abundant in the total RNA fraction of *ncb-1*. This may result from the strong nuclear retention of intron-retaining transcripts that are neither processed nor exported to the cytoplasm. These observations may indicate that the absence of CBs alters stress-induced changes in the processing and nuclear retention of transcripts, which may contribute to the reduced hypoxia tolerance of *ncb-1* (Górka et al., 2022).

### Involvement of Cajal bodies in nuclear RNA retention

Our results indicate that hypoxia leads to a pronounced redistribution, including selective retention, of poly(A^+^) transcripts between the nucleus and the cytoplasm, and that the presence of functional CBs affects the dynamics of this process, both at the stage of sequestration and at the stage of transcript release into the cytoplasm. During hypoxia, an increase in the nuclear poly(A^+^) RNA signal was observed with a concomitant decrease in its cytoplasmic level, whereas after reoxygenation a partial return to the distribution characteristic of control conditions occurred. This may indicate active sequestration of transcripts in the nuclear compartment rather than merely reduced export resulting from a global slowdown of metabolism. This phenomenon was previously described in preliminary form in root cells of *A. thaliana*, lupin and onion subjected to hypoxia (Niedojadło et al., 2014, 2016). In animal cells, stress-induced reorganization of nuclear speckles has likewise been linked to changes in the splicing of stress-responsive transcripts (Sung et al., 2023).

Nuclear RNA retention is a relatively recently recognized level of post-transcriptional gene regulation. In plants, its importance has been demonstrated both during development and in response to environmental factors (Göhring et al., 2014; Taliansky et al., 2023) and has also been documented in mammalian tissues (Bahar Halpern et al., 2015). Our results extend this concept by indicating that CBs may participate in regulating the selective retention and distribution of specific transcripts. Notably, despite the strong accumulation of poly(A^+^) RNA in nuclei observed by *in situ* methods, transcriptomic analysis did not indicate uniform, global retention of all transcripts. Instead, we identified transcripts at different stages of maturation under all conditions examined, including those containing retained introns (IR) and with differing IRX values in the nuclear and total RNA fractions. Although most strongly retained introns were shared between the genotypes (83.7%), about 8% were retained only in *ncb-1* and a similar proportion only in WT (Supplementary Table S3), indicating that CBs may modulate the retention of specific rather than random introns. This interpretation is supported by earlier observations showing that the lack of coilin in the *ncb-1* mutant leads to retention of introns 2 and 3 in RPB1 mRNA (Górka et al., 2022), whereas in larch microsporocytes CBs selectively retain mRNAs of Sm proteins containing specific retained introns (Rudzka et al., 2022). Introns retained exclusively in *ncb-1* may represent introns whose efficient splicing is supported by CBs. These results also show that the absence of properly functioning CBs does not abolish nuclear RNA accumulation but substantially alters the range of transcripts subject to this process. Their efficient splicing in WT during stress may be facilitated by CBs, potentially through the maintenance of a functional U2 snRNP complex. This is supported by our RIP-qPCR results showing higher binding of U2.2 and U2.3 snRNAs to the U2B″ protein in WT than in *ncb-1*, suggesting more efficient formation of functional U2 snRNP complexes in the presence of CBs.

It is also known that the nuclear localization of mRNA is frequently associated with the presence of retained introns (Lee and Bailey-Serres 2019). To resolve the heterogeneity of intron retention, we analysed separately introns with low (IRX < 10%) and high (IRX > 50%) retention, which revealed clearly distinct behaviour of these two groups in nuclear RNA. Efficiently spliced introns constitute by far the largest group, and during hypoxia they accounted for over 70% (WT) and 80% (*ncb-1*) of the introns detected in the nuclear fraction. In WT, this proportion increased two- to threefold relative to normoxia, whereas in *ncb-1*, in which it was already high under control conditions, the increase was only moderate. Given the concomitant accumulation of poly(A^+^) RNA in the nucleus (Fig. 1), this suggests that during hypoxia WT nuclei accumulate largely spliced transcripts, including those of genes involved in protein transport and ribosome biogenesis, and that CBs affect the dynamics of this process. The marked increase in splicing efficiency in WT under hypoxic conditions is likely related to the presence of CB as a site for snRNP maturation.

Nevertheless, intron-retaining transcripts did not disappear from the nucleus during hypoxia. Strongly retained introns still accounted for about 2–17% of the introns detected in the nuclear fraction of WT, and the last intron remained the least efficiently spliced under all conditions (Fig. 9). Their share, however, was markedly lower than under normoxia, and mean nuclear IRX decreased by 11–18 percentage points in WT but by only 2–3 points in *ncb-1* (Fig. 7C, D). During hypoxia, the nuclear pools of the two genotypes therefore became similar in their proportion of intron-retaining transcripts. The difference lay mainly in the dynamics: in WT, a large pool of incompletely spliced transcripts present under normoxia was depleted during stress, whereas in *ncb-1* this pool was small from the outset and changed little. In total RNA, by contrast, hypoxia slightly increased intron retention to a similar extent in both genotypes (Fig. 7A, B), suggesting that some intron-retaining transcripts may leave the nucleus or accumulate outside it during stress independently of CBs. Thus, although intron retention contributes to the nuclear RNA pool during hypoxia, it is not the main feature of the transcripts that accumulate in the nucleus under these conditions.

The second group comprises strongly retained introns, which were most abundant in the nuclear RNA of WT plants under normoxia and whose proportion decreased sharply during hypoxia (Fig. 9C). In *ncb-1*, strongly retained introns were less frequent under control conditions, and their proportion changed little during hypoxia (Fig. 9D). Genes containing such introns in nuclear RNA were enriched in GO terms related to abiotic stress, including hypoxia (Fig. S7A, B). Consistently, strongly retained introns were retained only under control conditions and spliced under stress, and this group was dominated by WT-specific introns (Supplementary Table S3). In addition, among the transcripts relatively enriched in the nuclear fraction under normoxia and shifted towards the total RNA pool during hypoxia (100 in WT and 127 in *ncb-1*; Fig. 4A, B), over 30 were associated with stress responses, including 5 related to hypoxia (Supplementary Table S2). In this way, plants may prepare in advance for potential stress. Only isolated examples of this have been described in the literature. One such example is the transcription factor PtOCR2-L in black cottonwood (*Populus trichocarpa*), whose mRNA contains retained introns 4 and 5 under control conditions (Filichkin et al., 2018). A similar phenomenon has been described in *A. thaliana* for the *CCA1* (CIRCADIAN CLOCK ASSOCIATED 1) transcript, which under physiological conditions is present in the nucleus as an intron-retaining form, whereas heat stress promotes removal of the intron and increases the proportion of the functional, fully spliced isoform (James et al., 2012; Filichkin et al., 2015). Our results indicate that this type of regulation may be more widespread. Importantly, the last intron was more frequently strongly retained than the first one (Fig. 9C, D), which may indicate a particular role of intron position in priming plants for abiotic stress. We observed differences between genotypes, in WT, the transition from normoxia to hypoxia was accompanied by a strong, more than threefold decrease in the number of introns in this group, whereas in *ncb-1* this effect was much weaker or absent. This suggests that functional CBs may promote the efficient removal of strongly retained, including terminal, introns during stress. At the same time, the level of such introns in the cytoplasm remained relatively stable, with only a slight increase observed in *ncb-1* during hypoxia. This may indicate that regulation of this group of transcripts through intron retention and removal occurs primarily at the nuclear level, since the amount of such introns that could be degraded or stored in the cytoplasm does not change. This indicates that transcriptome-level stress priming is considerably more active in WT plants and may be an important mechanism regulating abiotic stress tolerance.

### Sequence and positional determinants of selective intron retention

Retained introns in *A. thaliana* differed from efficiently spliced ones in a consistent set of features. Their weaker polypyrimidine tracts, more frequent non-GT–AG termini and a subset of severely weakened 5′ splice sites agree with previous reports identifying weak splice signals as a hallmark of retained introns (Mao et al., 2017; Dvinge 2018). The strongest discriminating feature, however, was the reduced compositional contrast between the intron and its flanking exons. Retained introns were more exon-like, which may weaken the UA-rich intron–exon contrast that the plant spliceosome relies on to define intron boundaries. The enriched motifs point to possible regulators. A-rich motifs may resemble binding sites of some SR proteins or A-rich elements near plant polyadenylation signals (Loke et al. 2005). CU-rich motifs may recruit PTB proteins, which compete with U2AF65 for the polypyrimidine tract and can thereby limit 3′ splice site recognition (Singh et al., 1995; Saulière et al., 2006). Two of the three *Arabidopsis* PTB homologs regulate intron retention transcriptome-wide (Rühl et al., 2012; Simpson et al., 2014), making them plausible candidates. Conversely, the U- and A-rich motifs enriched in efficiently spliced introns are consistent with the role of UA-rich intronic splicing enhancers in plants (Simpson et al., 2004).

We observed preferential retention of the first, second and last introns, whereas the compositional differences between retained and efficiently spliced introns were the same in all positional classes (Fig. S9); intron sequence alone therefore does not explain this positional pattern. Length asymmetry may play a role: globally, retained introns are longer than spliced introns (median 133 nt vs. 103 nt), but for the first intron from the 5′ end the relationship is reversed (152 nt vs. 184 nt), as it is in single-intron genes (178 nt vs. 296 nt).

At the 5′ end, shorter introns have less space for dispersed, redundant ISE motifs (including UA-rich sequences essential for intron definition in plants) (Mao et al., 2014; Rose et al., 2008), which we also identified in efficiently removed introns. Less space means a lower probability of containing motifs that bind splicing factors, which favors retention (Monteuuis et al., 2019). A similar pattern (weaker splice sites and depletion of TTTT-containing ISE motifs, together with enrichment of AG/GA-rich intronic splicing silencer (ISS) motifs in retained introns) was also reported in an independent comparative analysis of *Arabidopsis* introns (Mao et al., 2017).

For last introns, additional factors may contribute. First, for kinetic reasons: more than half of introns remain unspliced when Pol RNA II has moved more than 1 kb beyond the 3′ splice site, and the rate of co-transcriptional splicing in plants is considerably slower than in yeast (Jia et al., 2020); in the case of the last intron, this extends transcription time at the expense of the time available for splicing before termination and polyadenylation. Second, for sequence-related reasons: retained introns in *Arabidopsis* are enriched in intronic splicing silencers (ISSs), including GA-rich motifs (Mao et al., 2017), which is also supported functionally by experimental evidence showing that GAAG repeats act as an intronic splicing regulator in the SCL33 gene (Thomas et al., 2012). Our results refine this picture by revealing enrichment of A-rich motifs, a CG-containing motif and CU repeats. This is noteworthy because earlier comparative analyses had instead indicated generally lower GC content in retained introns than in constitutively spliced introns in *Arabidopsis* (Marquez et al., 2012); our finding of a specific CG-containing motif in retained introns may therefore represent a novel regulatory element. A longer intron statistically has more space for the accumulation of such elements, so kinetic and sequence factors may act synergistically to increase the likelihood of retention. These explanations remain hypothetical and require experimental verification.

Collectively, our results indicate that the function of Cajal bodies in the hypoxia response relies on several interconnected mechanisms. CBs promote the assembly of functional splicing complexes, including the association of U2 snRNA with the U2B″ protein into the U2 snRNP complex, and the reduction of this association in *ncb-1* may affect splicing efficiency in the mutant. Independently, CBs contribute to the transcriptional activation of hypoxia-responsive genes, whose induction in total RNA is stronger in WT than in *ncb-1*. With respect to nuclear RNA, the function of CBs manifests differently depending on the condition: during hypoxia, the presence of CBs is associated with a strong shift of the nuclear RNA pool towards efficiently spliced transcripts, whereas under normoxia WT nuclei contain a distinct pool of strongly retained, incompletely spliced transcripts of stress-responsive genes, which is depleted upon stress. Together, these data indicate that Cajal bodies coordinate snRNP maturation, transcriptional activation of stress genes, and the selective nuclear retention and redistribution of transcripts, suggesting that they are an important element of the molecular strategy by which plants adapt to, and recover from, oxygen deprivation.

## Supporting information

Differential expression of genes associated with systemic, nuclear, and spinal hypoxia

The table lists transcripts classified as switching between nuclear and total RNA fractions

Sequence and structural features of retained versus spliced introns

## Notes

### Competing Interest Statement

The authors have declared no competing interest.

### Funding

- National Science Centre: grant no. 2016/21/B/ NZ3/00367

