## Additional data for "Cajal bodies shape the nuclear transcriptome, intron retention and transcripts redistribution in *Arabidopsis* roots under hypoxia"

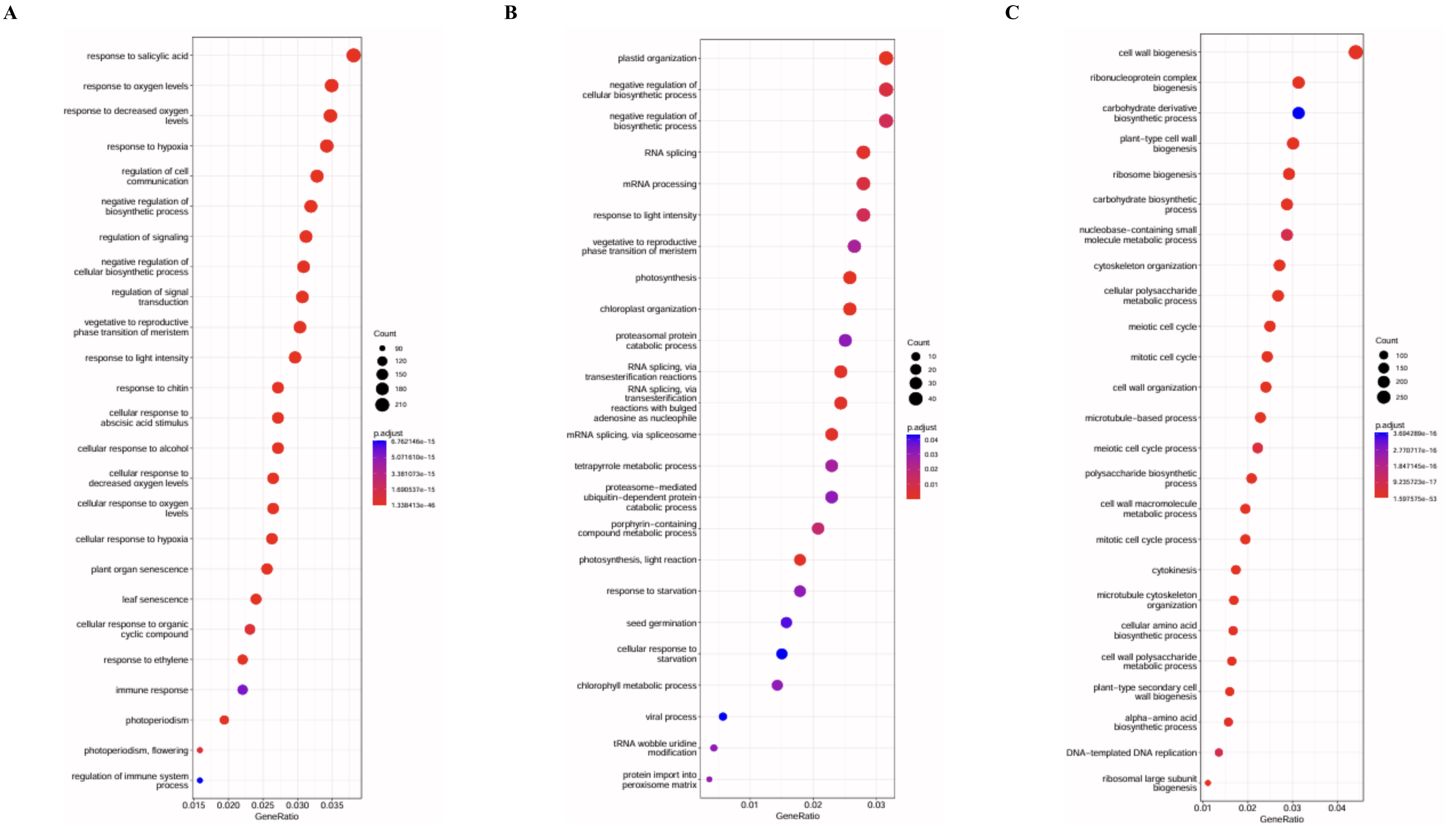


**Supplementary Figure S1 A-C.** Gene Ontology enrichment analysis of Biological Process terms for genes differentially expressed during hypoxia in total RNA. **(A)** Genes upregulated during hypoxia relative to control conditions in both WT and *ncb-1*. **(B)** Genes upregulated during hypoxia specifically in WT. **(C)** Genes downregulated during hypoxia in both WT and *ncb-1*. In all panels, the y-axis shows enriched GO terms and the x-axis shows GeneRatio, defined as the proportion of genes in the analyzed gene set assigned to each GO term. Dot size represents the number of genes associated with each term (Count), and dot color represents the adjusted *P* value (p.adjust).

D E F

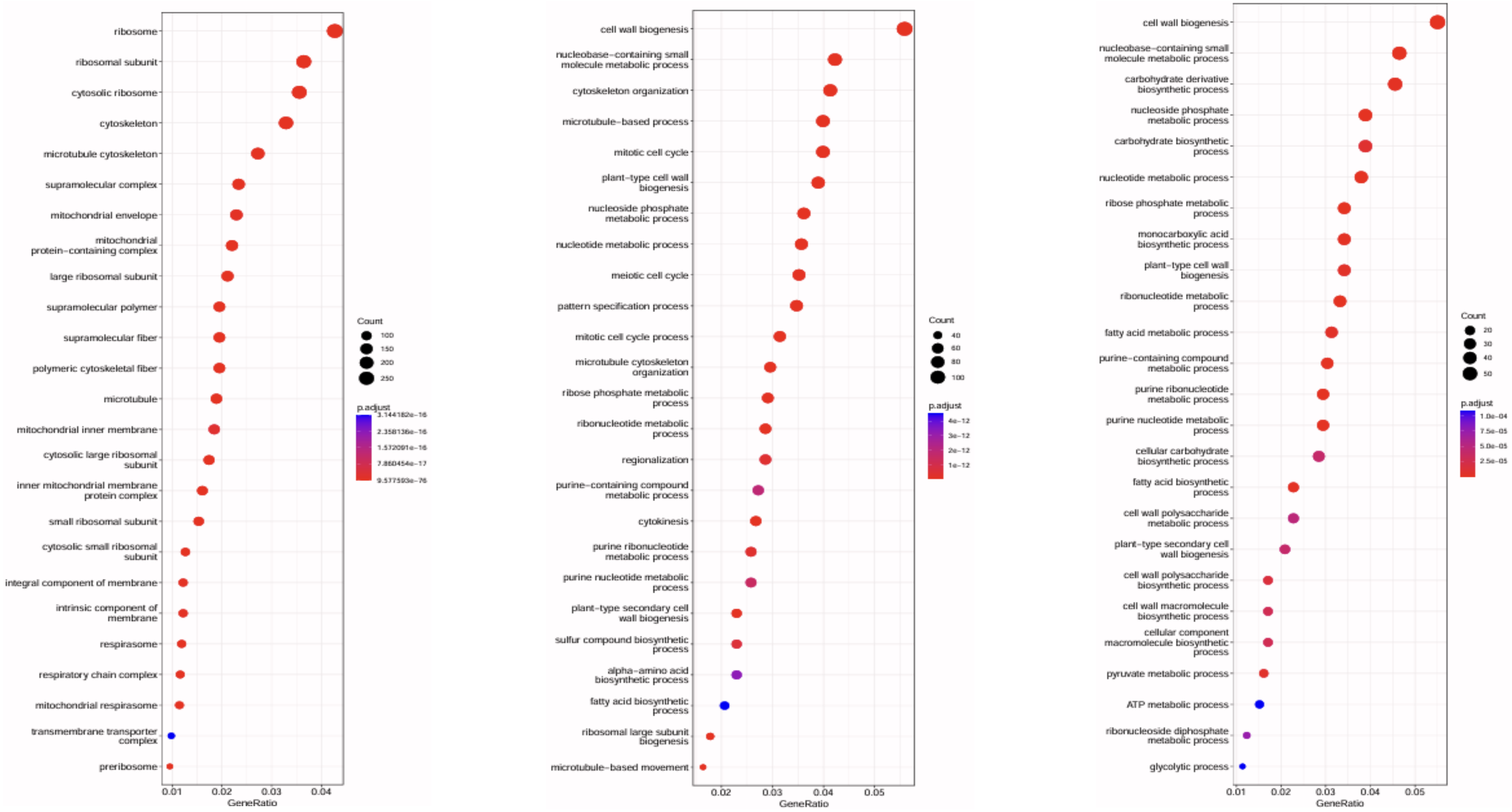


**Supplementary Figure S1 D-F.** Gene Ontology enrichment analysis of Cellular Component and Biological Process terms for genes differentially expressed during hypoxia and reoxygenation in total RNA. **(D)** Cellular Component enrichment analysis of genes downregulated during hypoxia relative to control conditions in both WT and *ncb-1*. **(E)** Biological Process enrichment analysis of genes upregulated during reoxygenation relative to hypoxia in both WT and *ncb-1*. **(F)** Biological Process enrichment analysis of genes upregulated during reoxygenation specifically in WT. Across all panels, enriched GO terms are displayed on the y-axis, while the x-axis represents GeneRatio, corresponding to the fraction of genes in the analyzed gene set assigned to each GO term. Dot size indicates the number of genes associated with a given term (Count), whereas dot color denotes the adjusted P value (p.adjust).

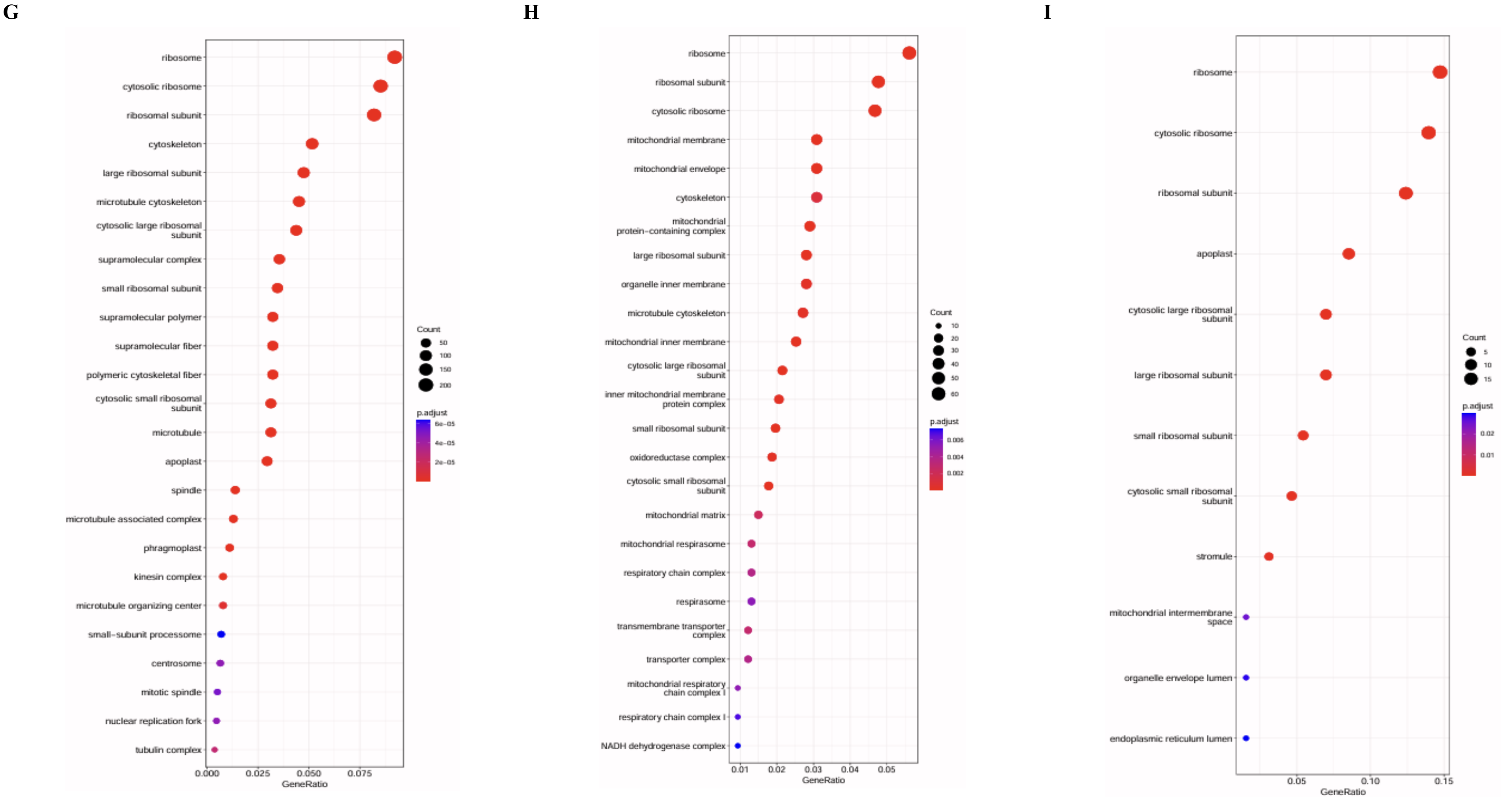


**Supplementary Figure S1 G-I.** Gene Ontology enrichment analysis of Cellular Component terms for genes upregulated in total RNA during reoxygenation. **(G)** Genes upregulated during reoxygenation relative to hypoxia in both WT and *ncb-1*. **(H)** Genes upregulated during reoxygenation specifically in WT. **(I)** Genes upregulated during reoxygenation specifically in *ncb-1*. For each panel, enriched GO terms are shown along the y-axis and GeneRatio is plotted on the x-axis, representing the proportion of genes from the analyzed gene set assigned to the respective GO term. The number of genes contributing to each term is reflected by dot size (Count), and the adjusted P value (p.adjust) is indicated by dot color.

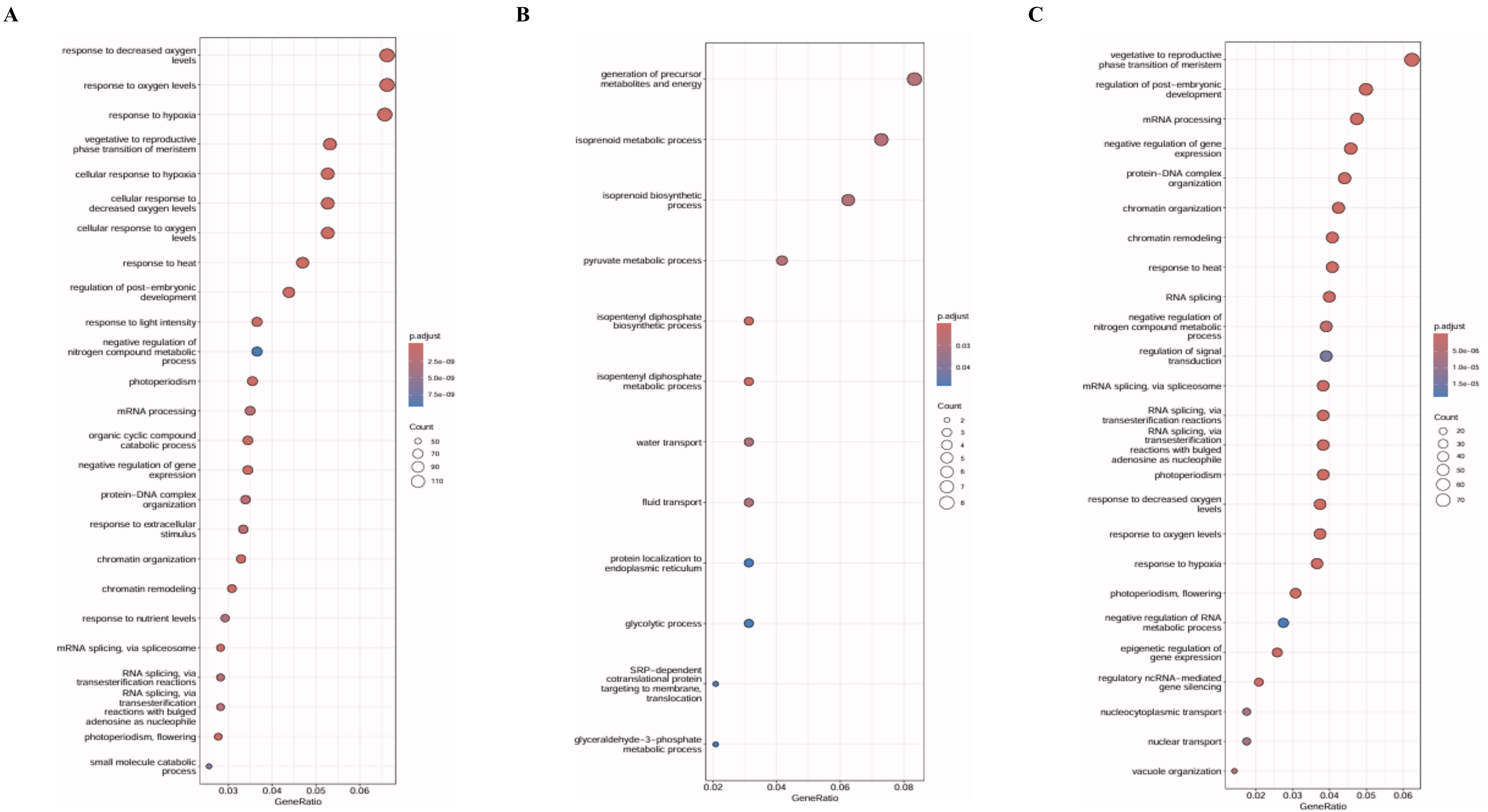


**Supplementary Figure S2 A-C.** Gene Ontology enrichment analysis of Biological Process terms for genes upregulated during hypoxia in the nuclear RNA fraction. **(A)** Genes upregulated during hypoxia relative to control conditions in both WT and *ncb-1*. **(B)** Genes upregulated during hypoxia specifically in WT. **(C)** Genes upregulated during hypoxia specifically in *ncb-1*. In all panels, the y-axis lists the enriched GO terms, whereas the x-axis corresponds to GeneRatio, defined as the proportion of genes in the analyzed set associated with each GO term. Dot size reflects the number of genes assigned to each term (Count), while the color scale represents the adjusted P value (p.adjust).

**D** **E** **F**

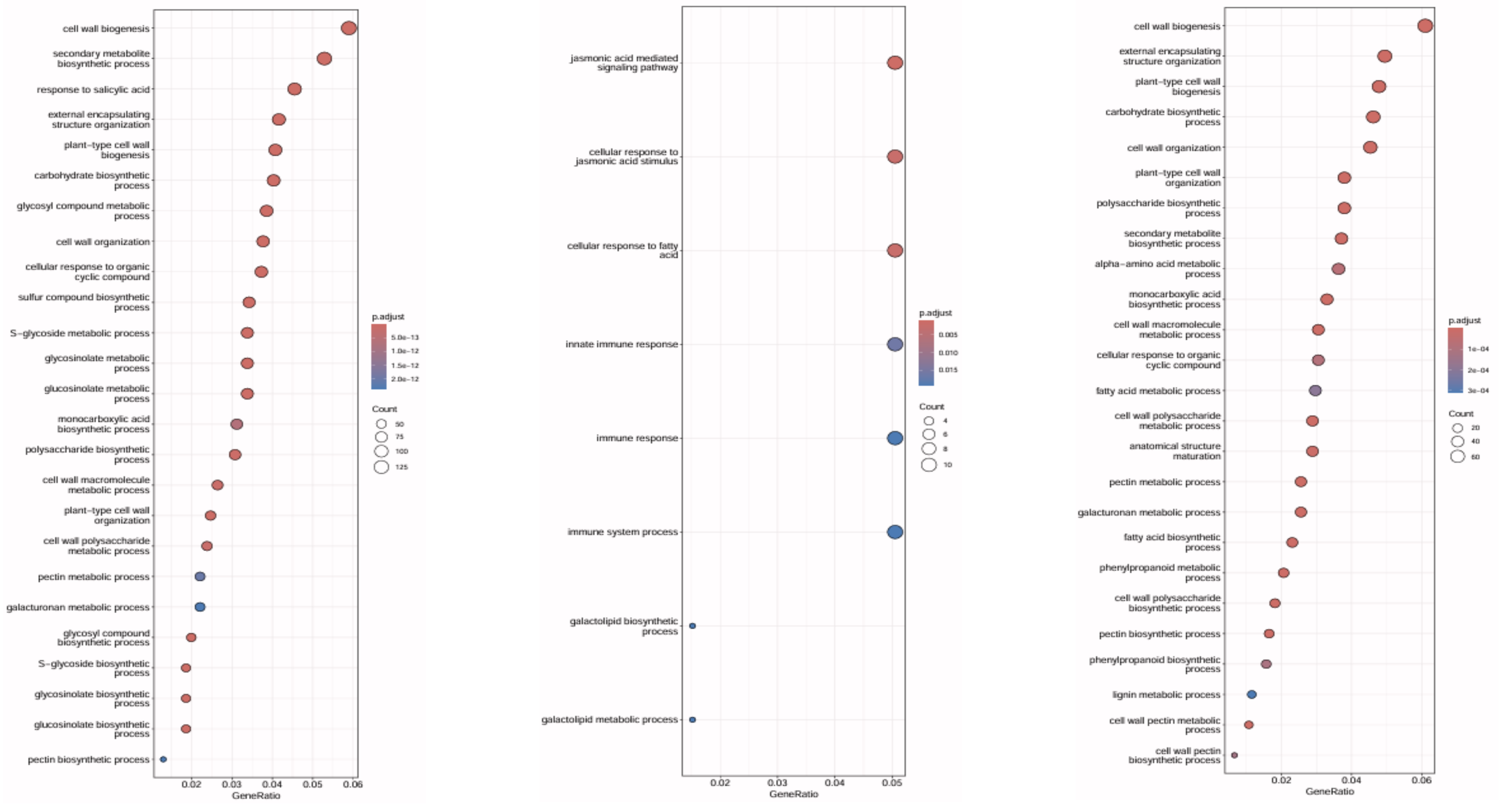


**Supplementary Figure S2 D-F.** Gene Ontology enrichment analysis of Biological Process terms for genes downregulated during hypoxia in the nuclear RNA fraction. **(D)** Genes downregulated during hypoxia relative to control conditions in both WT and *ncb-1*. **(E)** Genes downregulated during hypoxia specifically in WT. **(F)** Genes downregulated during hypoxia specifically in *ncb-1*. Enriched GO terms are presented on the y-axis in each panel, with GeneRatio shown on the x-axis as the fraction of genes within the analyzed gene set linked to a particular GO term. The size of the dots corresponds to the number of associated genes (Count), and their color reflects the adjusted P value (p.adjust).

**A**

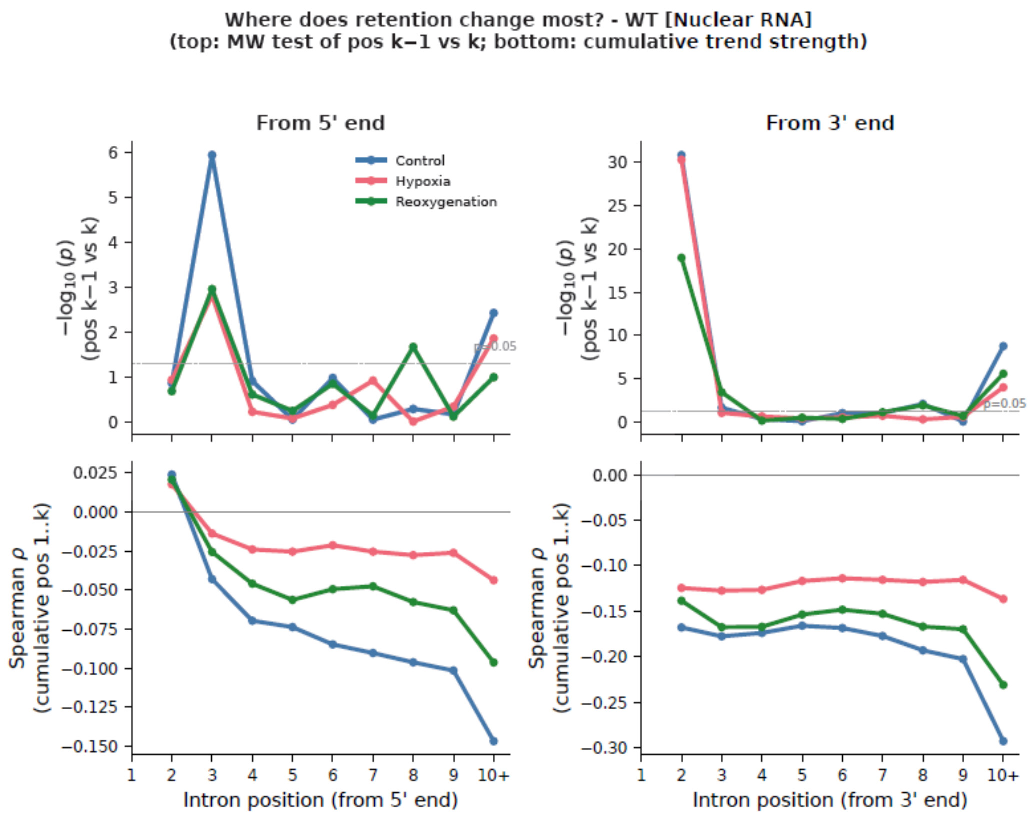


**B**

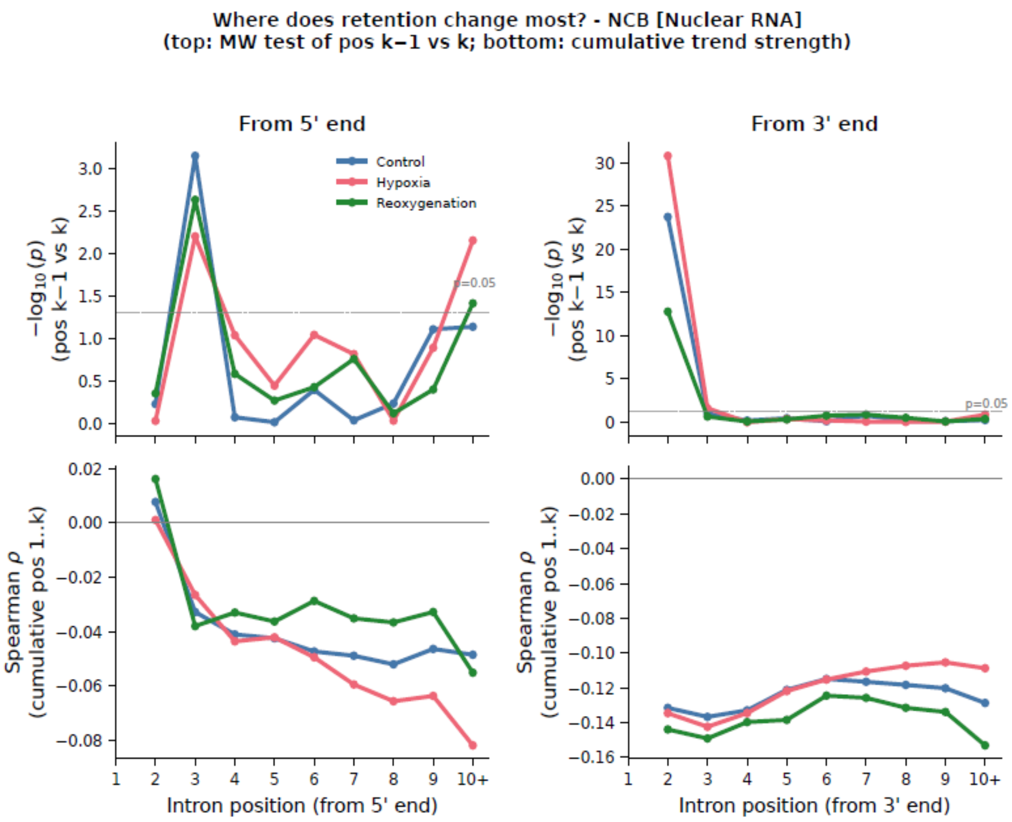


**Supplementary Figure S3 A, B. Position-dependent changes in intron retention in WT and *ncb-1* nuclear RNA.(A)** Positional analysis of intron retention in WT nuclear RNA under control, hypoxia, and reoxygenation conditions. **(B)** Corresponding analysis for the *ncb-1* nuclear RNA fraction. Intron positions were analyzed separately from the **5′ end** (left panels) and the **3′ end** (right panels) of the gene. Upper panels show the statistical significance of positional differences in intron retention, expressed as −log10($P$) from Mann–Whitney tests comparing each intron position with the preceding position ($k$ − 1 vs. $k$). The horizontal line indicates the significance threshold of $P = 0.05$. Lower panels show the cumulative Spearman correlation coefficient (ρ) between intron position and intron retention, providing a measure of the direction and strength of the positional trend. Blue, red, and green lines represent control, hypoxia, and reoxygenation conditions, respectively.

**A** **B**

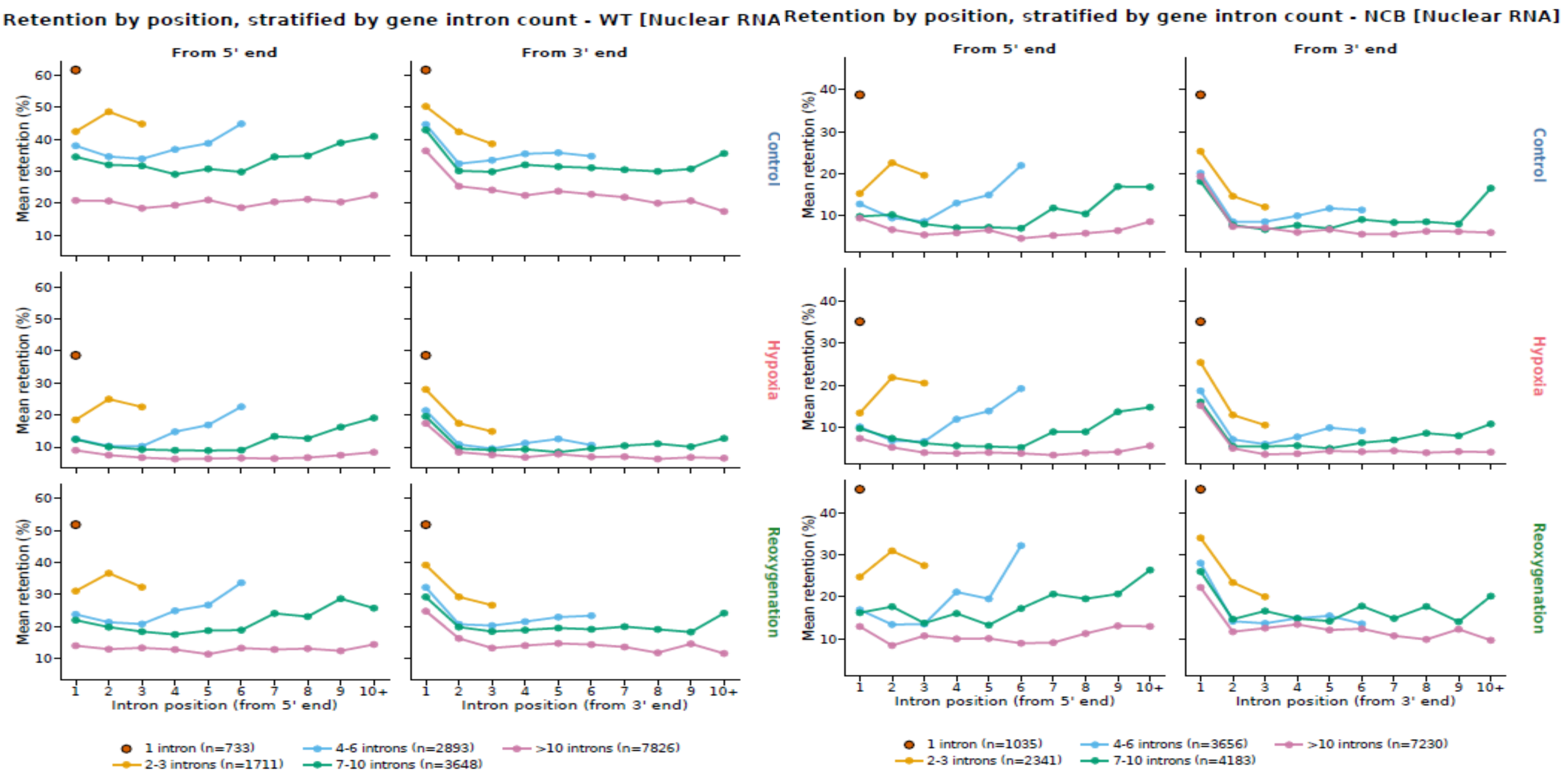


**Supplementary Figure S4 A, B. (A)** Mean intron retention according to intron position in the WT nuclear RNA fraction under control, hypoxia, and reoxygenation conditions. **(B)** Corresponding analysis for the *ncb-1* nuclear RNA fraction. Introns were stratified according to the total number of introns per gene (1 intron, 2–3 introns, 4–6 introns, 7–10 introns, and >10 introns). Intron positions were analyzed separately relative to the **5′ end** (left panels) and **3′ end** (right panels) of the gene. Rows represent control (top), hypoxia (middle), and reoxygenation (bottom) conditions. The y-axis shows mean intron retention (%), and the x-axis indicates intron position counted from the respective gene end.

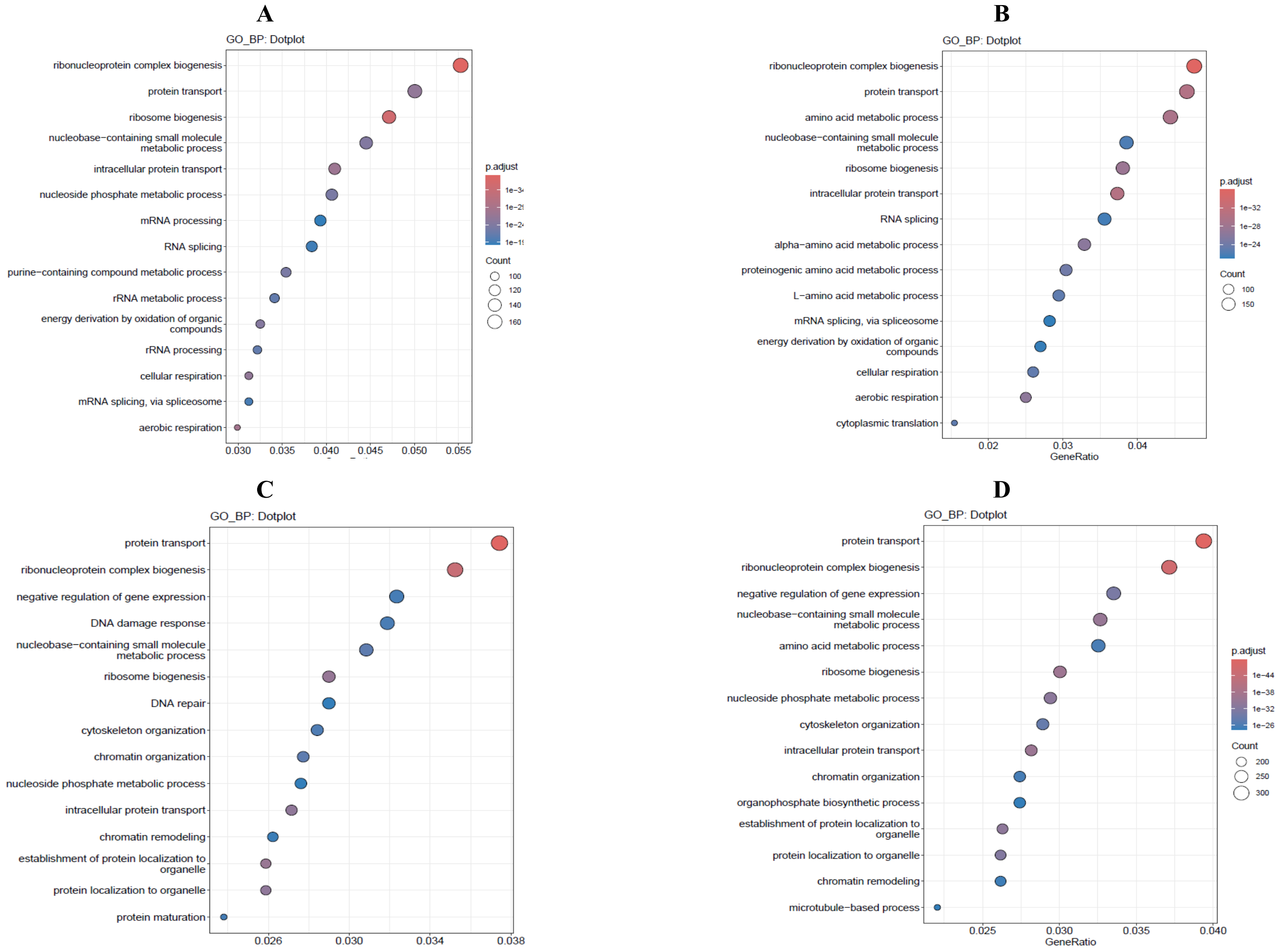


**Supplementary Figure S5 A-D.** Gene Ontology Biological Process (GO:BP) enrichment analysis of genes containing introns with an intron retention index (IRX) below 10% in nuclear and total RNA fractions of WT and *ncb-1 A. thaliana*. (**A**) GO:BP enrichment analysis for genes containing introns with IRX < 10% in the nuclear RNA fraction of WT *A. thaliana*. **(B)** Corresponding GO:BP enrichment analysis for the nuclear RNA fraction of the *ncb-1* mutant. **(C)** GO:BP enrichment analysis for genes containing introns with IRX < 10% in the total RNA fraction of WT *A. thaliana*. **(D)** Corresponding GO:BP enrichment analysis for the total RNA fraction of the *ncb-1* mutant. In all panels, enriched GO terms are displayed on the y-axis, whereas the x-axis represents GeneRatio, defined as the proportion of genes from the analyzed gene set assigned to each GO term. Dot size reflects the number of genes associated with a given term (Count), while dot color indicates the adjusted P value (p.adjust).

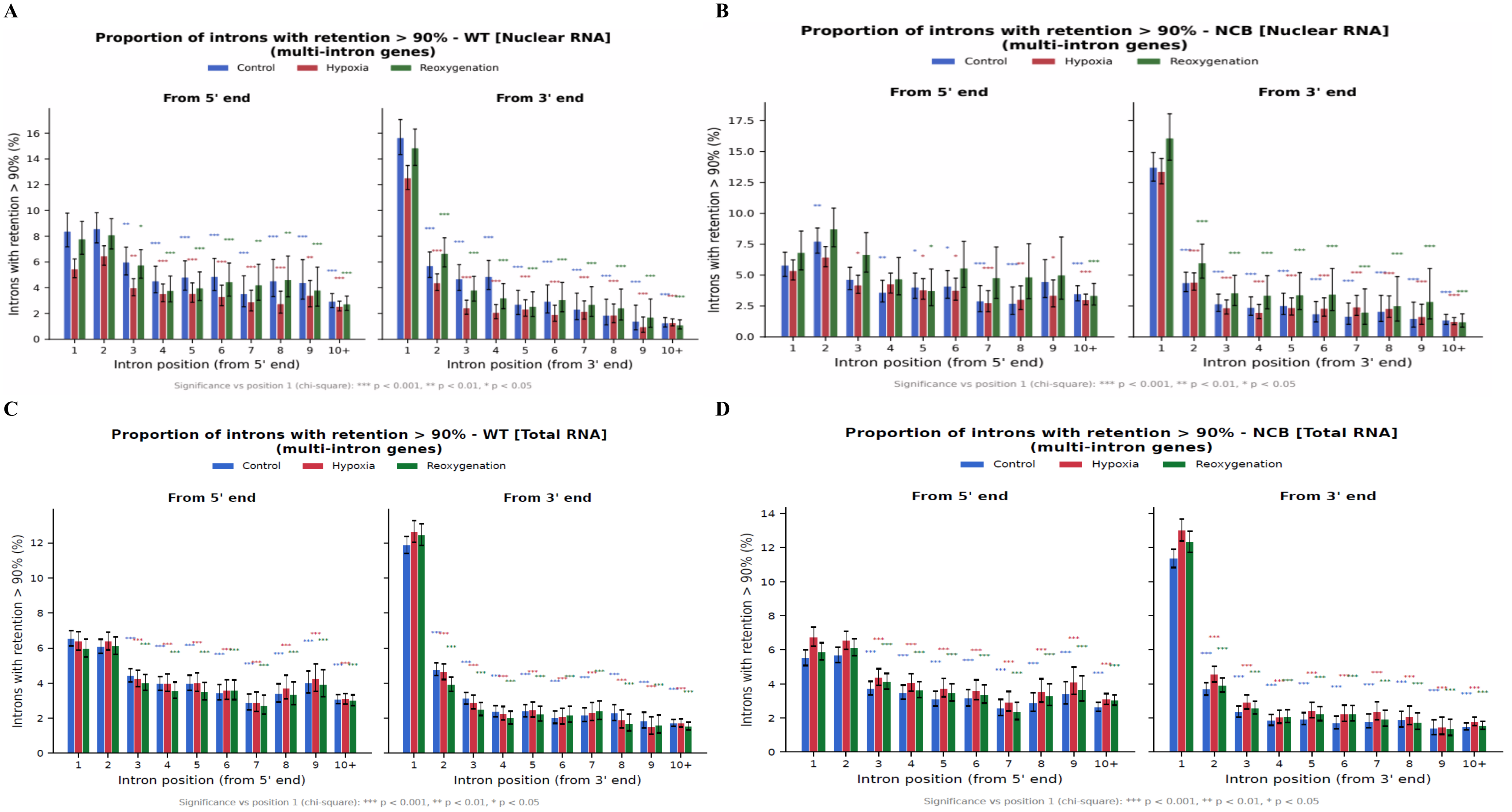


**Supplementary Figure S6 A-D.** Positional distribution of strongly retained introns (IRX > 90%) in multi-intron genes in nuclear and total RNA fractions of WT and *ncb-1 A. thaliana*. **(A)** Proportion of introns with IRX > 90% in the nuclear RNA fraction of WT *A. thaliana*. **(B)** Corresponding analysis for the nuclear RNA fraction of the *ncb-1* mutant. **(C)** Proportion of introns with IRX > 90% in the total RNA fraction of WT *A. thaliana*. **(D)** Corresponding analysis for the total RNA fraction of the *ncb-1* mutant. In each panel, intron positions are shown separately from the 5′ and 3′ ends of multi-intron genes. The x-axis indicates intron position counted from the respective gene end, whereas the y-axis represents the proportion of introns with IRX > 90%. Bars correspond to control, hypoxia, and reoxygenation conditions. Statistical significance relative to position 1 is indicated by asterisks as shown in the plots (chi-square test; ***P < 0.001, **P < 0.01, *P < 0.05).

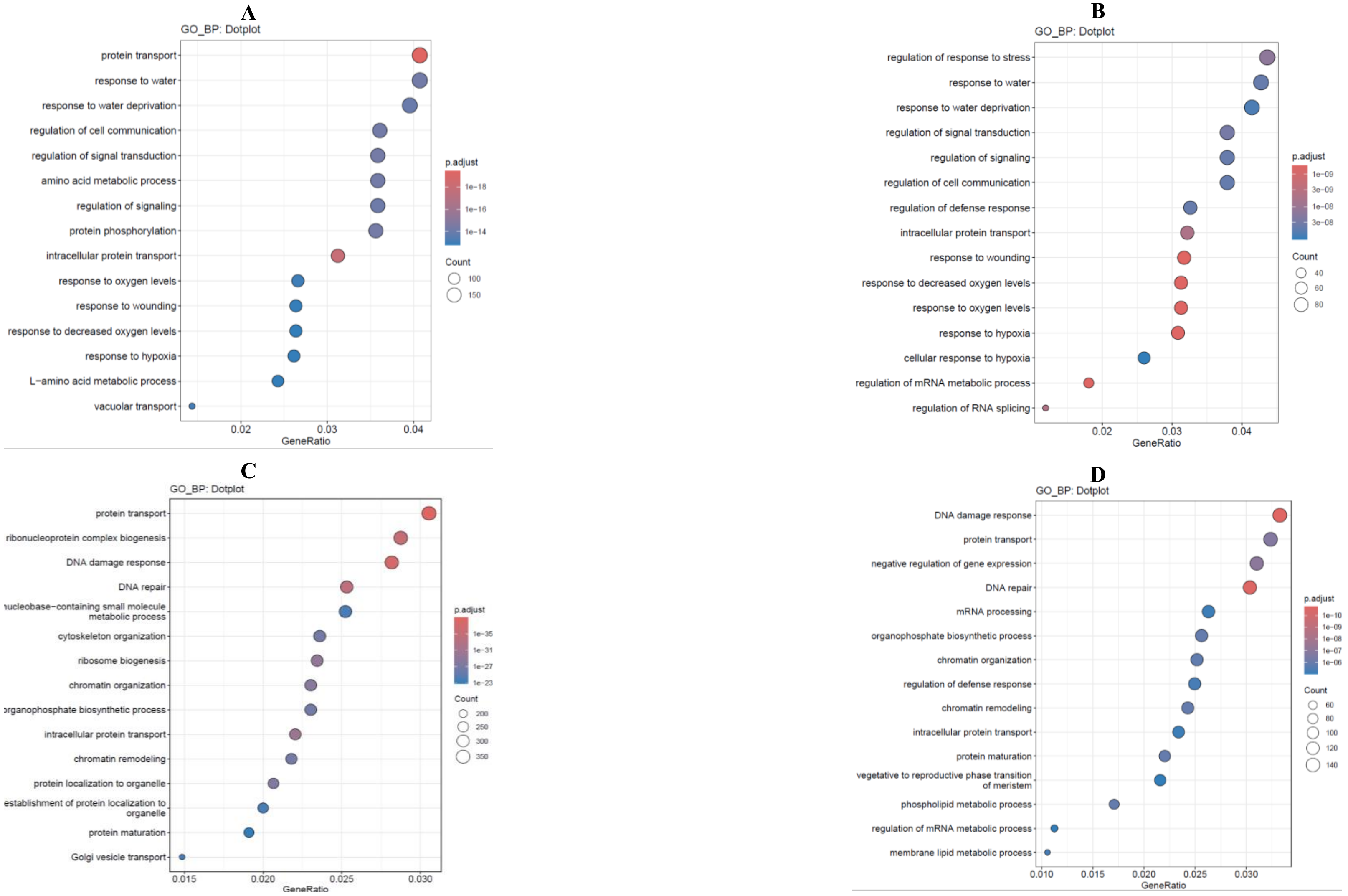


**Supplementary Figure S7 A-D.** Gene Ontology Biological Process (GO:BP) enrichment analysis of genes containing introns with an intron retention index (IRX) above 50% in nuclear and total RNA fractions of WT and *ncb-1 A. thaliana*. **(A)** GO:BP enrichment analysis for genes containing introns with IRX > 50% in the nuclear RNA fraction of WT *A. thaliana*. **(B)** Corresponding GO:BP enrichment analysis for the nuclear RNA fraction of the *ncb-1* mutant. **(C)** GO:BP enrichment analysis for genes containing introns with IRX > 50% in the total RNA fraction of WT *A. thaliana*. **(D)** Corresponding GO:BP enrichment analysis for the total RNA fraction of the *ncb-1* mutant. In all panels, enriched GO terms are displayed on the y-axis, whereas the x-axis represents GeneRatio, defined as the proportion of genes from the analyzed gene set assigned to each GO term. Dot size reflects the number of genes associated with a given term (Count), while dot color indicates the adjusted P value (p.adjust).

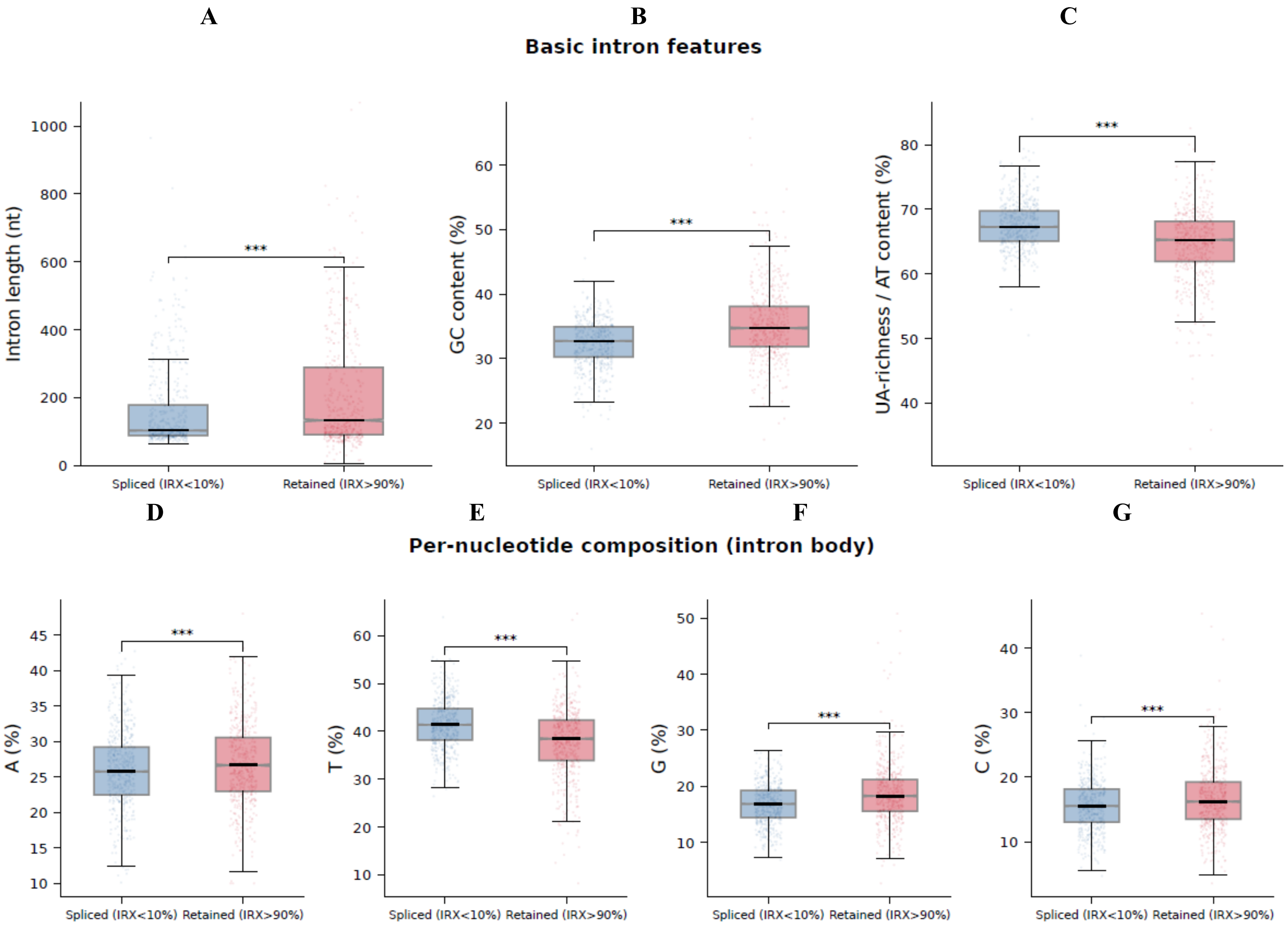


**Supplementary Figure S8 A-G.** Sequence and compositional features of low-retention and strongly retained introns. Introns with IRX < 10% (blue; low-retention/efficiently spliced) were compared with introns with IRX > 90% (red; strongly retained). **(A)** Intron length. **(B)** GC content. **(C)** UA-richness/AT content. **(D-G)** Relative nucleotide content of A, T, G, and C in the intron body, respectively. Box plots summarize the distributions of the two intron groups. **Statistical significance between low-retention and strongly retained introns was assessed using the Mann–Whitney U test; significance is indicated by asterisks (*P < 0.001).

H I

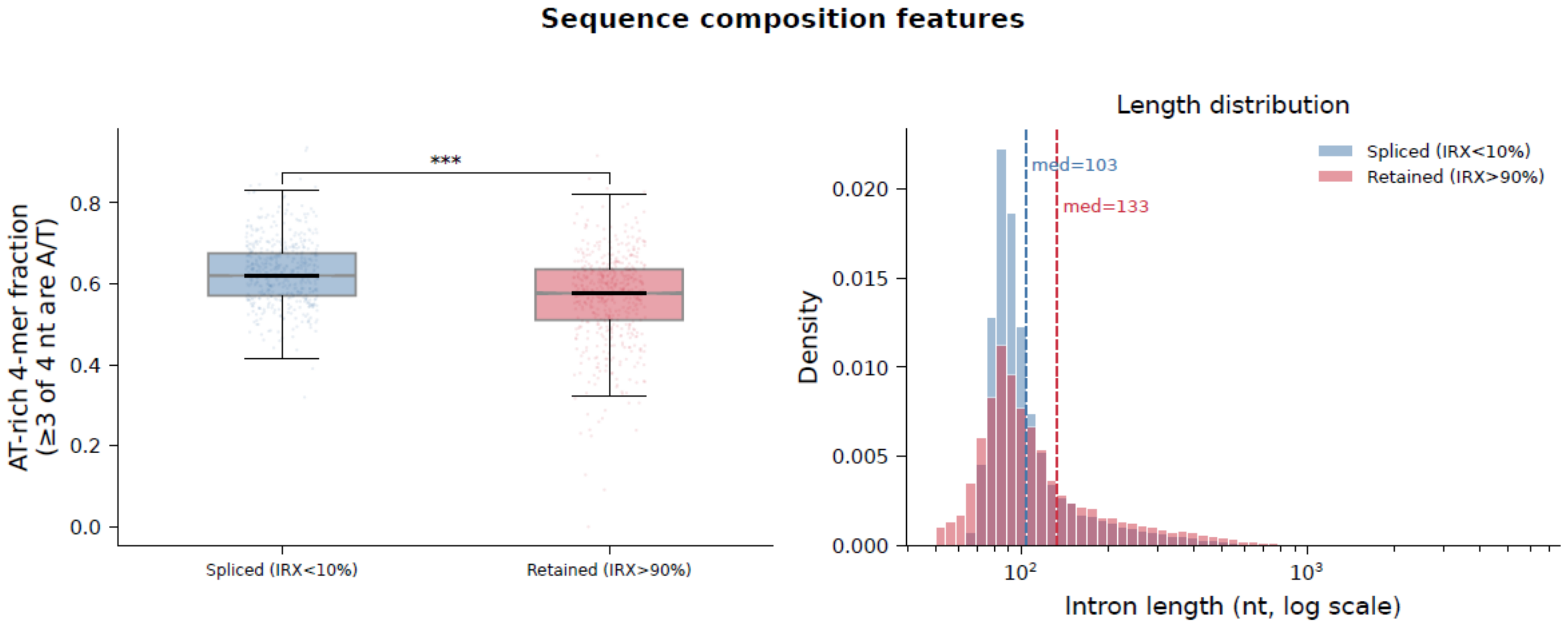


**Supplementary Figure S8H-I.** Additional sequence-composition and length features of low-retention and strongly retained introns. **(H)** Fraction of AT-rich 4-mers (at least 3 of 4 nucleotides are A/T) in introns with IRX < 10% and IRX > 90%. **(I)** Intron-length distributions for the same groups, displayed on a logarithmic x-axis; dashed vertical lines mark median lengths of 103 nt for IRX < 10% introns and 133 nt for IRX > 90% introns. Colors correspond to low-retention (blue) and strongly retained (red) introns. **Statistical significance was assessed using the Mann–Whitney U test; significance is indicated by asterisks (*P < 0.001).

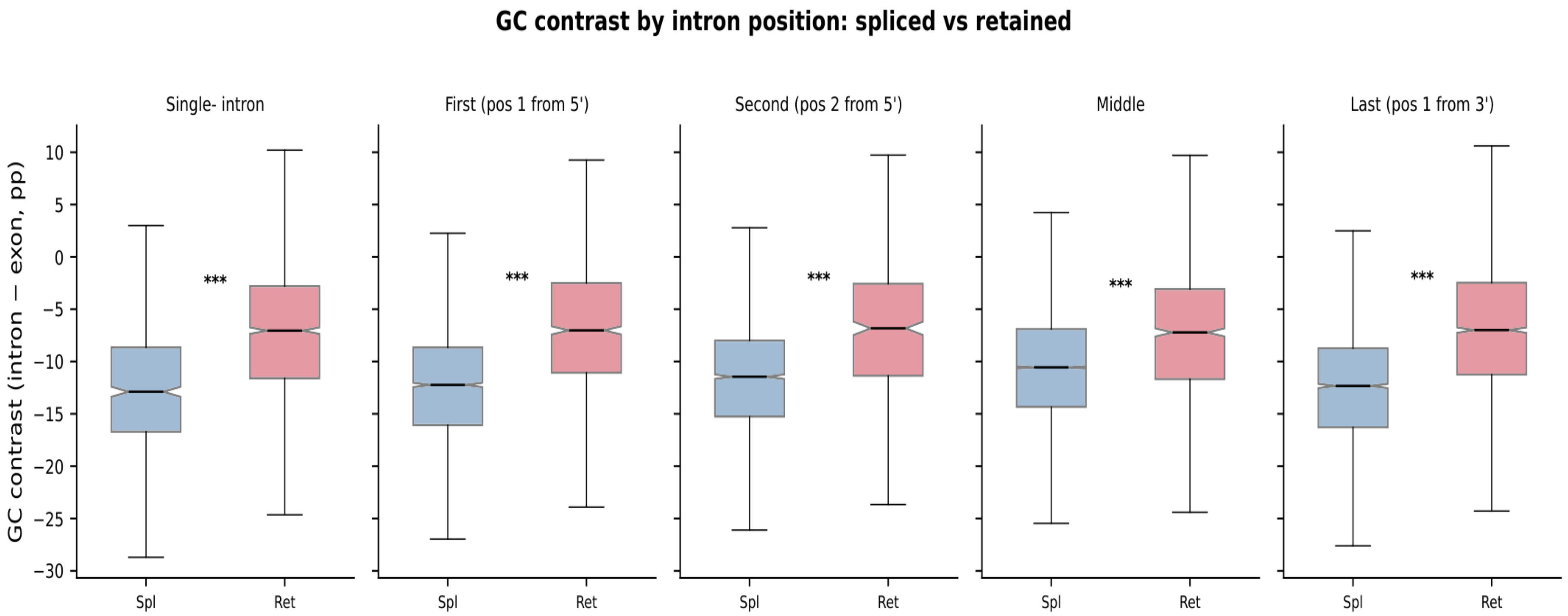


**Supplementary Figure S9. GC contrast between efficiently spliced and strongly retained introns across positional classes.** GC contrast between introns and their flanking exons (intron GC − exon GC, percentage points) was compared between efficiently spliced introns (Spl) and strongly retained introns (Ret) across five positional classes: single-intron, first (position 1 from the 5′ end), second (position 2 from the 5′ end), middle, and last (position 1 from the 3′ end) introns. Box plots show the distribution of GC contrast values within each positional class. Statistical significance between efficiently spliced and strongly retained introns is indicated by asterisks (***P < 0.001).

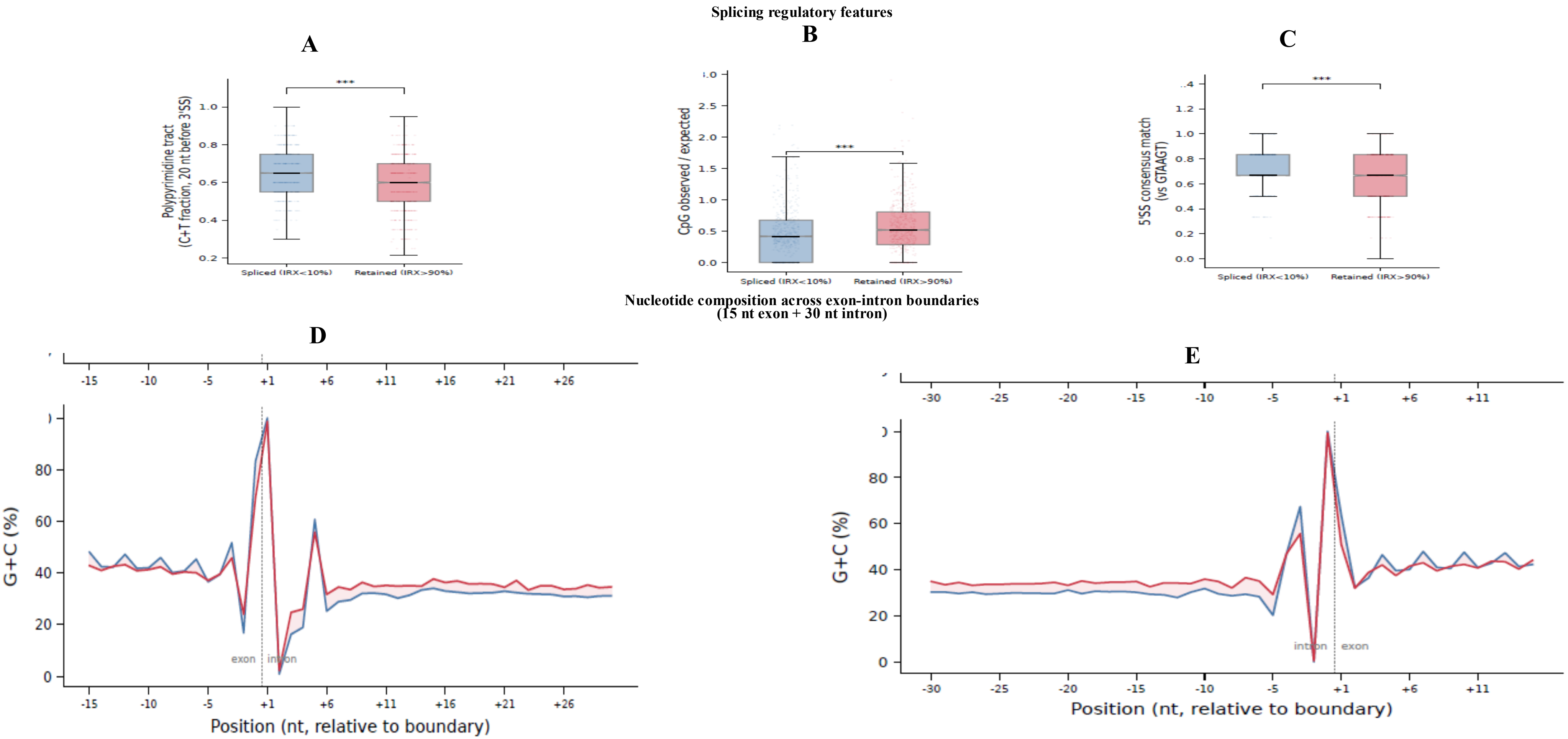


**Supplementary Figure S10A-E.**Splicing-regulatory sequence features and positional intron-retention patterns of low-retention and strongly retained introns. Introns with IRX < 10% (blue; low-retention/efficiently spliced) were compared with introns with IRX > 90% (red; strongly retained). **(A)** Fraction of pyrimidines (C+T) in the 20-nt polypyrimidine tract immediately upstream of the 3′ splice site. **(B)** CpG observed-to-expected ratio. **(C)** Match to the 5′ splice-site consensus sequence (GTAAGT). **(D-E)** G+C nucleotide-composition profiles across exon-intron boundaries. **(D)** G+C content around the 5′ splice site, covering the exon-to-intron transition. (E) G+C content around the 3′ splice site, covering the intron-to-exon transition.

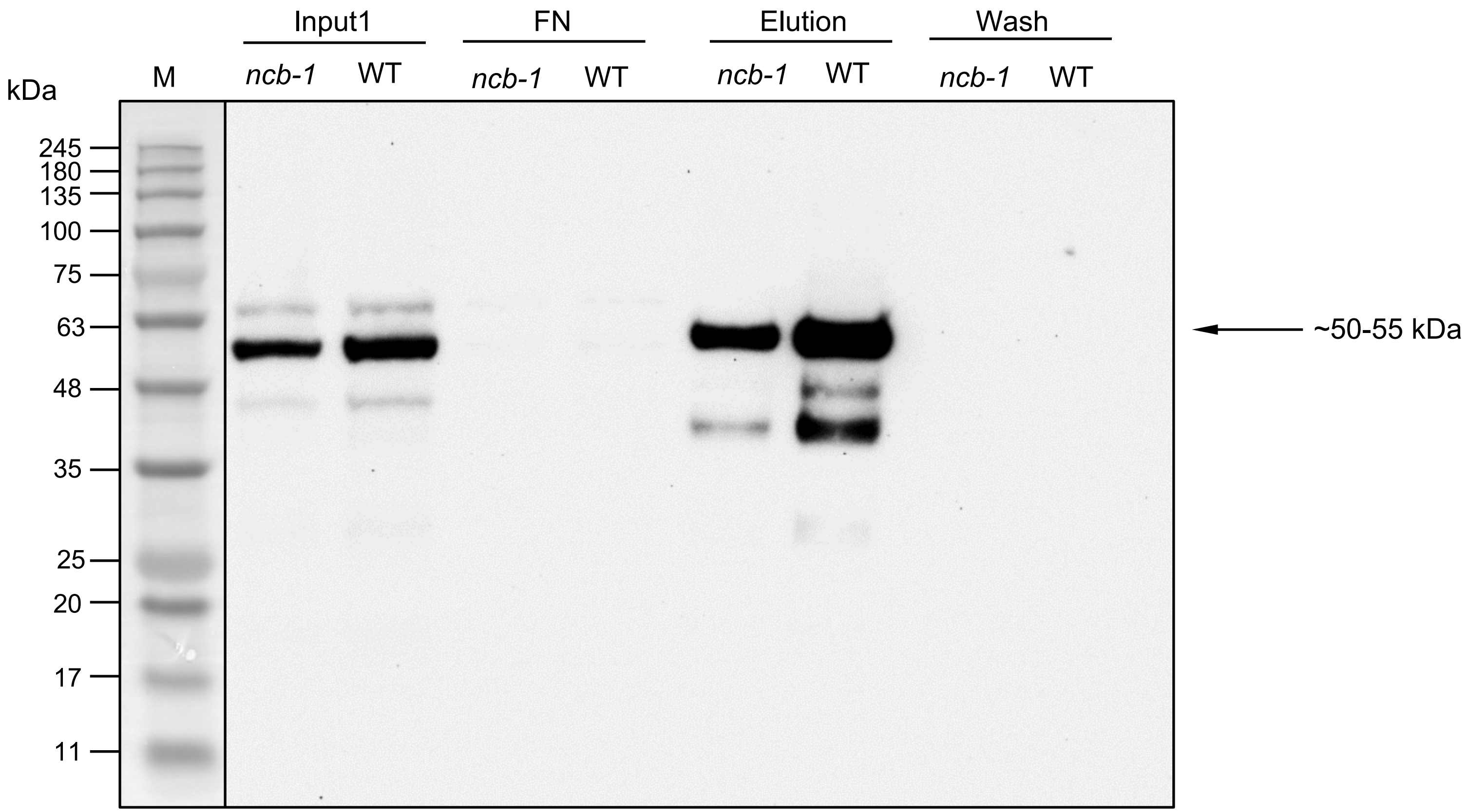


**SupplementaryFigureS11.** Immunoblot analysis of U2B-GFP-containing protein fractions from WT and *ncb-1 A. thaliana*. Input, flow-through/non-bound (FN), elution, and wash fractions obtained following GFP-Trap affinity purification were analyzed by immunoblotting using an anti-GFP-HRP antibody (Miltenyi Biotec; 1:1000). A prominent band at approximately 50–55 kDa, corresponding to the expected molecular weight of the U2B-GFP fusion protein**,** was detected in the input and elution fractions of both genotypes, with a visually stronger signal in WT than in *ncb-1*.Little or no U2B-GFP signal was detected in the wash or FN fractions. M indicates the molecular mass marker.
